# IL-6R blockade with tocilizumab disrupts pericyte– and tumor cell–driven IL-6/STAT3 signaling, enhancing docetaxel efficacy in ER+ breast cancer

**DOI:** 10.64898/2026.01.29.702661

**Authors:** Róża Przanowska, Jacky Gomez-Villa, Victoria J. Liu, Natalia Antonides-Jensen, Lavanya Visvabharathy, John C. Alverdy, Sonia L. Hernandez, Samantha S. Yee

## Abstract

Metastatic breast cancer is a global health concern with a persistently low five-year survival rate. Taxane microtubule stabilizers, including docetaxel (DTX), are the standard of care in various treatment protocols. DTX is used both as a single agent and in combination therapies, with a majority of ER+ breast cancer patients ultimately developing chemoresistance. The mechanisms contributing to chemoresistance involving the tumor microenvironment (TME) have not been fully elucidated. Specifically, the role of vascular cells within the TME, particularly pericytes, is understudied, and their role in promoting chemoresistance remains unknown. Inflammatory cytokines such as interleukin 6 (IL-6) are known to drive drug resistance via activation of the pro-survival JAK/STAT pathway. We found that DTX induced IL-6 secretion of pericytes by at least two-fold compared to vehicle-treated controls *in vitro*. All tested breast cancer cell lines expressed subunits of the IL-6 receptor (IL-6R) complex, indicating their capacity to respond to JAK/STAT signaling. Conditioned media from DTX-treated pericytes activated STAT3 in ER+ breast cancer cells to levels comparable to recombinant IL-6. Pharmacologic blockade of IL-6 signaling with the IL-6R inhibitor, tocilizumab, reduced DTX-induced STAT3 activation *in vitro.* Furthermore, combined treatment with tocilizumab and DTX synergistically suppressed the growth of zero-passage patient-derived ER+ breast cancer organoids expressing intact IL-6 signaling. Together, our findings suggest that combining DTX with tocilizumab may revert DTX-induced chemoresistance in ER+ breast cancer patients by inhibiting IL-6-mediated activation of the STAT3 pathway.

## Introduction

Breast cancer is the most prevalent cancer in women, excluding skin cancers. Within the USA, there are approximately 321,910 estimated new cases of breast cancer in women for 2026 ^1^. Luminal breast cancer, characterized by estrogen receptor positivity (ER+) and low human epidermal growth factor 2 (HER2), comprises 70% of all cases and generally has a favorable prognosis, with a 95.1% five-year survival rate^2^. Standard treatment involves surgery followed by endocrine-based pharmacologic therapy, but drug resistance—both acquired (40% of advanced cases) and intrinsic (over 50%)—remains a major challenge^3^. Consequently, 10% of ER+ patients develop distant metastases within ten years, higher than other breast cancer subtypes^4^. There was an estimation of approximately 170,000 women living with stage 4 metastatic breast cancer in 2025 within the USA^5^.

Microtubule targeting agents (MTAs) are chemotherapeutics used as single agents or in combination therapy for several cancer types and have interphase and mitotic effects depending on the concentration administered. Among the MTA agents, docetaxel (DTX, Taxotere) from the taxane class has been used as a first-line treatment for metastatic breast cancer^6^. DTX in combination therapy has greatly improved patient outcomes in both early-stage and metastatic breast cancer^7^. The efficacy of DTX has additionally been established in patients with ER+ metastatic breast cancer, with a series of randomized trials^8^.

Chemoresistance to taxanes occurs in about 30% of women with ER+ breast cancer, frequently leading to treatment failure^9,10^. Therapeutic resistance occurs when tumor cells overcome the toxic effects of chemotherapy, resulting in enhanced survival under drug-induced stress^11^. Drug resistance mechanisms, including the enhancement of DNA repair pathways, drug efflux transporters, disruption of cell cycle checkpoints, evasion of immune surveillance, and factors within the tumor microenvironment (TME), have a major influence on cancer progression and treatment resistance^12^. More specifically, the TME includes an abundance of cytokines as crucial messengers for crosstalk between different cell types^13^. Interleukin-6 (IL-6) is known to activate the Janus Kinase 2 (JAK2)/signal-transducer and activator of transcription 3 (STAT3) signaling pathway, resulting in chemoresistance^14–16^. This pathway has also been clinically validated as a therapeutic target for inflammation-related diseases^17,18^. Aberrations in this pathway have been strongly correlated with oncogenic processes such as proliferation, invasion, and metastasis in various malignancies, including breast cancer^19,20^.

Despite significant advances in breast cancer research, the TME persists as a central obstacle to the development of effective therapies for malignancies that carry a poor prognosis^21,22^. The TME is comprised of the internal and external environment of tumor cells as they replicate, invade adjacent tissues, and metastasize to secondary sites, with the environment being composed of unique cell types and crucial components, including immune cells, microvasculature, the extracellular matrix, fibroblasts, and cytokines, by all cells in the TME, such as interleukins^23^. This environment is rich in nutrients, dynamic, and is often favorable for tumor survival. Several studies have focused on the effects of DTX on cancer cells and fibroblasts for the TME^24,25^. Another crucial component of the TME is the microvasculature, which consists of pericytes and vascular smooth muscle cells (perivascular), and endothelial cells^26,27^, in which the tumor vasculature can support tumor progression^28,29^.

Pericytes have garnered significant attention in recent research over the past couple of decades, particularly in cancer. It is well established that pericytes remodel and stabilize the vasculature to ensure proper blood flow^30^. They also participate in blood vessel growth^31^. Pericytes are embedded between the basement membrane and endothelial cells and are part of the structure of human capillaries^32,33^. In cancer, they have been implicated in facilitating the metastatic process by allowing tumor cell intra- and extravasation^34–36^. They also play a role in immune tolerance^37^ and drug tolerance in cancer cells^38^. Pericytes are present in all solid tumors on most tumor vessels^39^, and are closely related to fibroblasts in the context of the breast^40^ and lung tissues^41^.

Furthermore, pericytes are not polarized cells and can secrete factors both into the surrounding tissue and directly into the bloodstream^42^. The pericyte secretome, deemed as “pericrine signaling,” is recognized as a key driver of tumor progression^34^. Even under basal conditions, the pericyte secretome is enriched in growth factors, cytokines/chemokines, extracellular matrix components, enzymes and proteases, and regulatory molecules, including IL-6, insulin-like Growth Factor 2 (IGF-2)^43^ and transforming growth factor (TGF)-β^44,45^, all of which can profoundly affect chemoresistance in tumor cells by activating unwanted inflammatory signaling pathways^46^. In colorectal cancer liver metastasis, tumor pericyte-secreted Wnt5a activates the STAT3 signaling pathway, promoting tumor progression^47^. Recent studies in thyroid and breast cancer, respectively, have shown that pericrine signaling contributes to anti-tumor efficacy^48^, tumor growth^43^, metastasis^49^, and regulates breast cancer dormancy^50^. There is a paucity of data on the effects of DTX on pericytes, particularly on its cytokine secretome^45,51–54^. Therefore, we asked whether pericytes have a bystander and/or active role on the cancer cells and disease outcomes, especially following chemotherapeutic (DTX) treatment.

We tested the hypothesis that elevated IL-6 in the secretome of DTX-treated pericytes contributes to chemoresistance through the JAK/STAT pathway in ER+ breast cancer using both *in vitro* and *ex vivo* models. Therefore, the aim of this study was to interrogate the direct effects of the pericyte secretome on ER+ breast cancer *in vitro* and *ex vivo*.

## Results

### Docetaxel induces IL-6 secretion in pericytes

Docetaxel (DTX) is a common chemotherapeutic used to treat breast cancer, but its effects on pericyte production of cytokines are unknown. We utilized a literature-based concentration of 10 nM DTX for 24-hour treatment to maintain consistency with published clinical findings^55,56^. 10 nM DTX represents a pharmacologically and clinically relevant concentration to assess the cellular impact and mechanisms of DTX^57–60^. Approximately 70% of pericytes and 80% of cancer cells survived following 24-hour treatment (Supplementary Fig. 1).

Cytokine secretion of human brain vascular pericyte (HBVP) cells was assessed after a 24-hour DTX or vehicle exposure. IL-6 production was increased by about two-fold (*P* = 0.0245) (Fig. 1a). In contrast, there was no statistical difference for other cytokines that may promote metastasis, such as CXCL1 (C-X-C Motif Chemokine Ligand 1)^61^, CXCL5 (C-X-C Motif Chemokine Ligand 5)^62^, ST2 (suppression of tumorigenicity 2)^63^, and GDF-15 (Growth Differentiation Factor 15)^64^ (Fig. 1a). IL-6 plays a critical role in tumor microenvironment (TME) interactions, enhancing the proliferation, survival, and metastasis of tumor cells through inflammatory pathways^65,66^, providing the rationale for focusing further on IL-6.

**Fig. 1:**
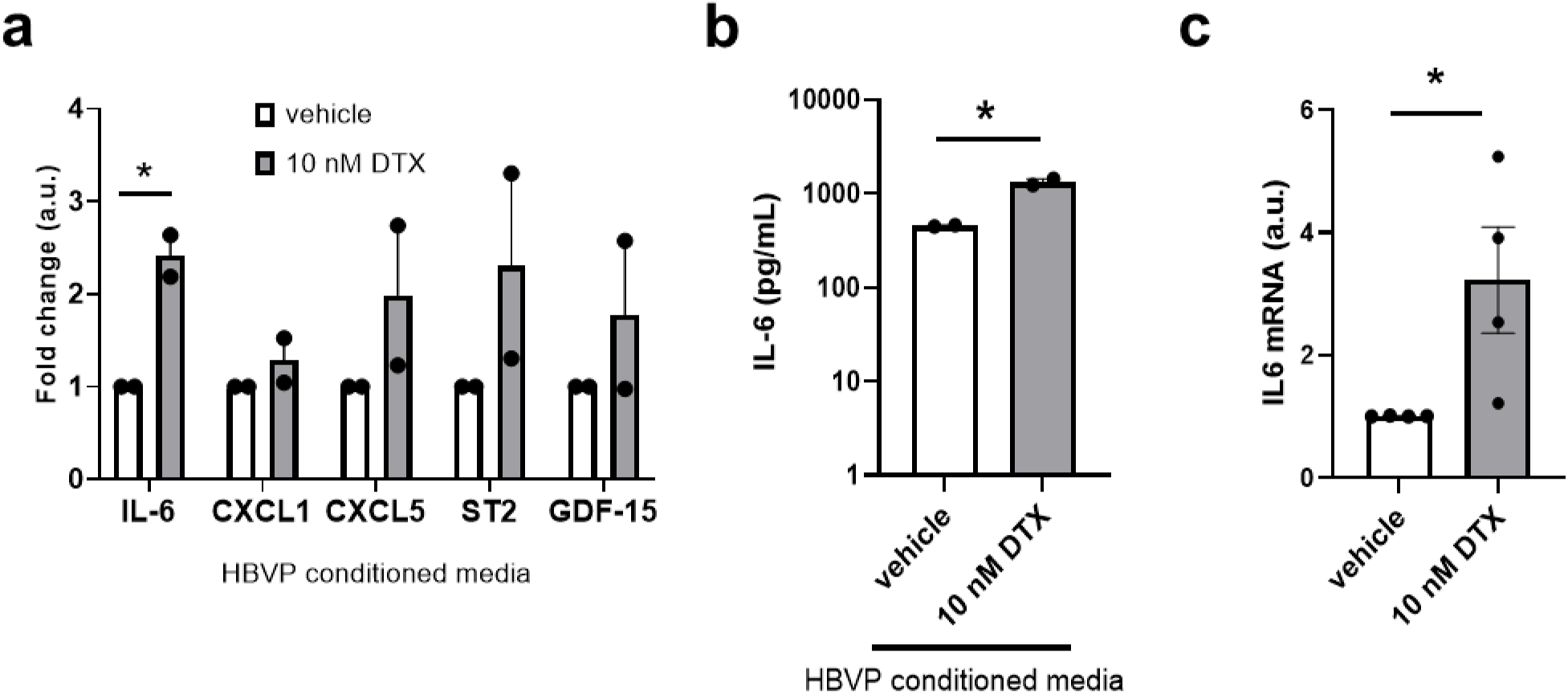
Docetaxel enhances IL-6 production and mRNA in pericytes. **a** Cytokine expression. Fold change of cytokine array data highlighting IL-6, CXCL1, CXCL5, ST2, and GDF-15 from HBVP conditioned media. a.u. denotes arbitrary units. Data is representative of n = 2 independent experiments with technical duplicates. Mean ± s.e.m. shown. **b** IL-6 (pg/mL) from HBVP conditioned media quantified by IL-6 ELISA. Data is representative of n = 2 independent experiments with technical duplicates. Mean ± s.e.m. shown. **c** *IL6* transcript levels in reference to GAPDH. a.u. denotes arbitrary units. Data is representative of n = 4 independent experiments with technical duplicates. Mean ± s.e.m. shown. *P* values were calculated by using an unpaired two-tailed t-test for **a, b, and c**. * *P* = 0.0245 (**a**), * *P* = 0.0134 (**b**), and * *P* = 0.0431 (**c**). * *P* ≤ 0.05. HBVP = human brain vascular pericytes. DTX = docetaxel.

To validate the cytokine array findings, IL-6 secretion was further assessed. Physiologically, reported levels from a meta-analysis of homeostatic IL-6 from the blood of healthy donors range from approximately 0–43.5 pg/mL^67^. During cancer development and treatment, exposure to infection^68^, or inflammatory diseases, IL-6 serum levels are elevated to the lower ng/mL range^69,70^. Supernatant from DTX-treated HBVP cells showed almost three-fold induction of IL-6 (Fig. 1b; *P* = 0.0134). DTX-treated HBVP cells had a greater than three-fold increase of *IL6* mRNA (Fig. 1c; *P* = 0.0431). Collectively, these results demonstrated that 10 nM DTX exposure for 24 hours induces IL-6 expression and secretion from the HBVP cells, which are then free to interact with other cells or with themselves.

### Basal IL-6, IL-6Ralpha, and IL-6Rbeta expression from pericytes, normal breast epithelial cell lines, and breast cancer cell lines

To assess autocrine and paracrine IL-6 signaling and establish its baseline capacity in the ER+ breast cancer TME^71–73^, we examined the expression of IL-6 and its receptor subunits in a cellular panel system consisting of HBVP cells, two non-tumorigenic breast epithelial cell lines (184A1 and MCF-10A), one triple-negative breast cancer (TNBC) cell line (MDA-MB-231), and three ER+ breast cancer cell lines (MCF-7, T-47D, and HCC1500) (Supplementary Table 1). ER+ breast cancer cell lines had the lowest levels of *IL6* mRNA expression, while the TNBC cell line and HBVP cells had the highest levels (Supplementary Fig. 2a). The HBVP cells secreted high levels of IL-6 compared to non-tumorigenic breast epithelial and ER+ breast cancer cell lines (*P* ≤ 0.0001), but not for the TNBC cell line (*P* = 0.8862) (Fig. 2a). The findings indicate that the HBVP cells have higher basal IL-6 secretion and higher levels of *IL6* mRNA expression than the ER+ breast cancer cell lines and non-tumorigenic breast epithelial cell lines. Therefore, under basal conditions, paracrine IL-6 signaling driven by pericytes may be more prominent than autocrine signaling from the ER+ breast cancer cell lines.

**Fig. 2:**
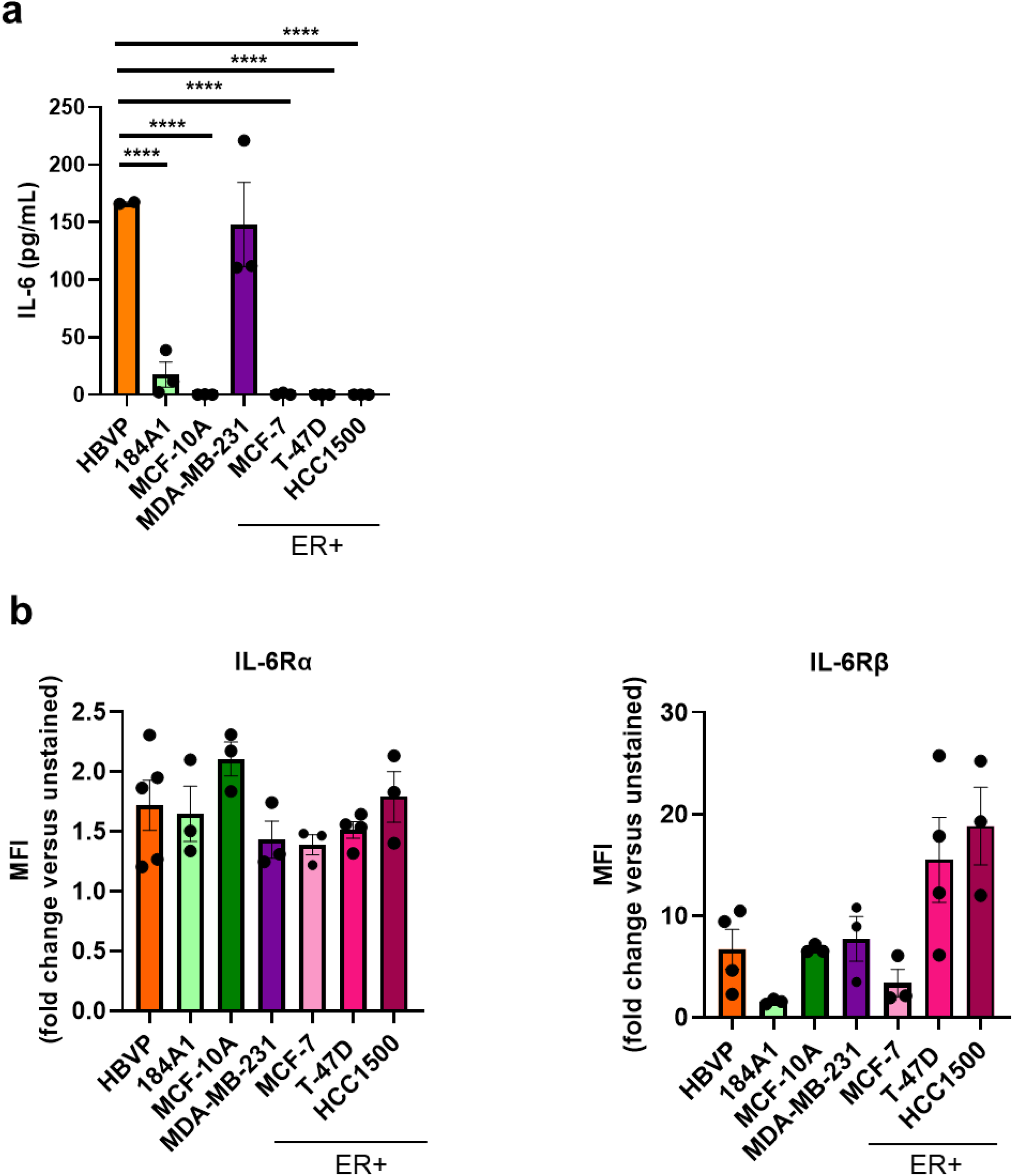
Pericytes secrete high levels of IL-6, while pericytes, normal breast epithelial cell lines, and breast cancer cell lines express IL-6Rα and IL-6Rβ. **a** Secreted IL-6 (pg/mL) from conditioned media of untreated cells and cell lines quantified by an IL-6 ELISA. Data is representative of n = 2 - 3 independent experiments. Mean ± s.e.m. shown. **b** MFI (fold change versus unstained) of IL-6Rα (left) and IL6Rβ (right) stained cells via flow cytometry. MFI = median fluorescence intensity. Data is representative of n = 3 – 5 independent experiments. Mean ± s.e.m. shown. a *P* values were calculated using an ordinary one-way ANOVA with a Dunnett’s multiple comparisons test compared to the HBVP cells. **** *P* ≤ 0.0001. HBVP = human brain vascular pericytes.

Next, we interrogated this cell line and the primary cell panel for the expression of the IL-6 receptor (IL-6R) complex, which binds IL-6 and activates the pro-survival JAK/STAT pathway^73,74^. The IL-6R complex consists of two subunits, alpha and beta, encoded by *IL6R* and *IL6ST* genes, respectively. All tested cell lines and primary cells express both *IL6R* and *IL6ST* mRNA, but at varying levels (Supplementary Fig. 2b). The MCF-10A, T-47D, and 184A1 cell lines had the highest levels of *IL6R* mRNA expression compared to the rest of the cells (Supplementary Fig. 2b left). All cells had comparable IL-6Rα protein levels at the cell surface (Fig. 2b left). The T-47D cell line had the highest levels of *IL6ST* mRNA expression, followed by the HCC1500 cell line, compared to the rest of the other cells tested (Supplementary Fig. 2b right). IL-6Rβ protein levels at the cell surface were higher in the T-47D cell line, with approximately 20-fold higher levels than the MCF-7 and 184A1 cell lines (Fig. 2b right). This observation is consistent with prior literature showing higher *IL6ST* expression in the T-47D cell line compared to the MCF-7 cells^75^. In summary, all cell types examined express IL-6Rα and IL-6Rβ, indicating that they are capable of responding to IL-6 and activating the downstream JAK/STAT signaling pathway that promotes cancer cell survival.

### IL-6 activates STAT3 in ER+ breast cancer cell lines

Because IL-6 promotes therapeutic resistance through activation of STAT3^76^, we evaluated basal and IL-6–stimulated STAT3 activation in our breast cancer cell lines. Basal STAT3 activation was assessed in untreated cells and cell lines cultured in their complete media (Fig. 3a). Constitutive STAT3 phosphorylation was found in HBVP cells and the TNBC cell line, whereas ER+ breast cancer cell lines had lower basal STAT3 activation (Fig. 3a). More specifically, the MDA-MB-231 cell line, and HBVP cells have high STAT3 activation, whereas the HCC1500 cell line has lower levels of STAT3 phosphorylation followed by the MCF-7, and T-47D cell lines which have minimal to non-detectable basal STAT3 activation.

**Fig. 3:**
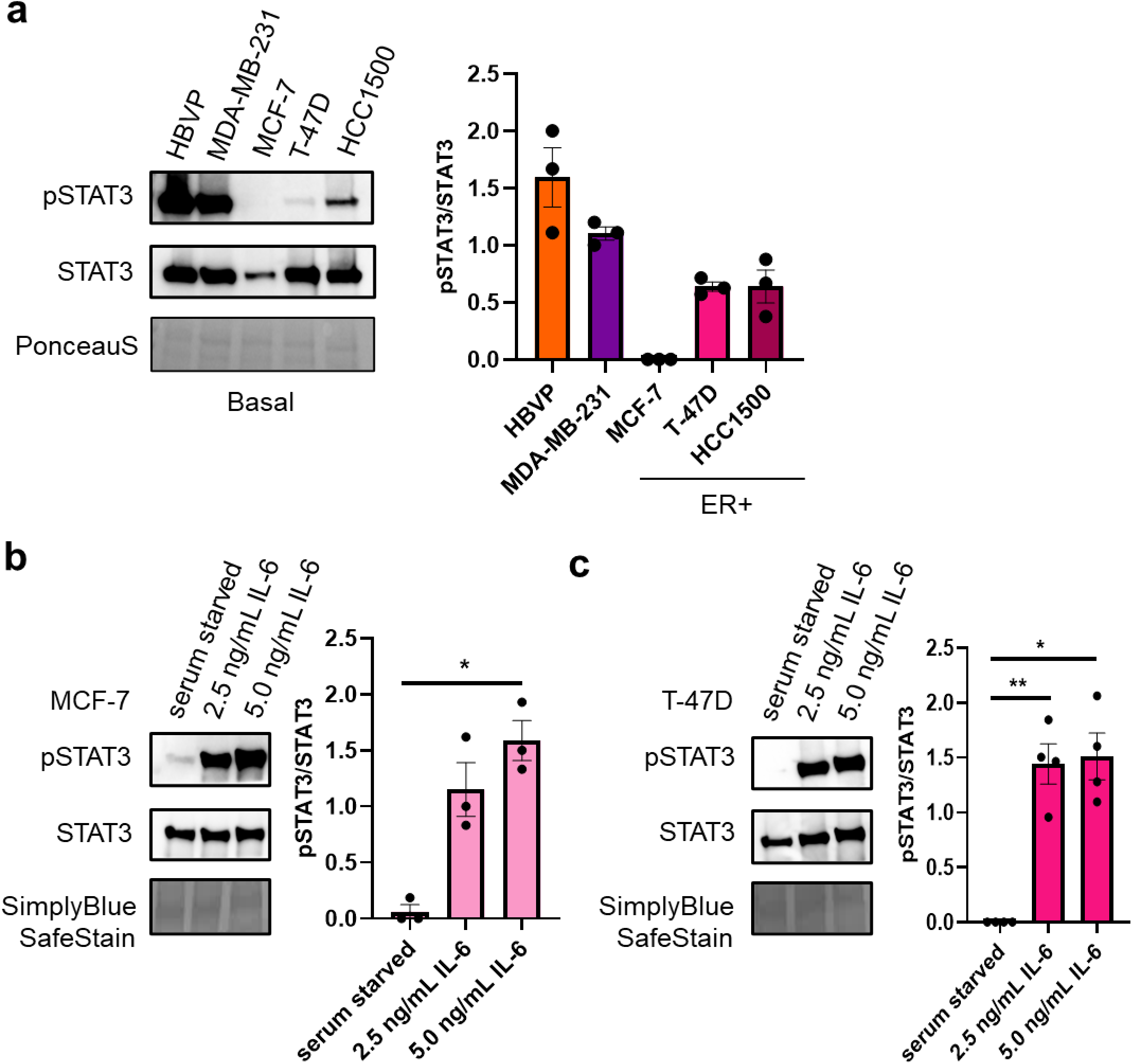
STAT3 (Tyr705) activation levels in pericytes and breast cancer cell lines, basally and following recombinant human IL-6 treatment. **a** Basal HBVP, MDA-MB-231, MCF-7, T-47D, and HCC1500 cells were lysed and subjected to immunoblotting using a pSTAT3 or STAT3 antibody. Fold change of pSTAT3 over total STAT3. Data is representative of n = 3 independent experiments. **b** MCF-7 and **c** T-47D cells were serum starved, then incubated with DMEM SFM (control media), 2.5 ng/mL IL-6 or 5.0 ng/mL IL-6 alone as indicated for 24 hours. Cells were then lysed and subjected to immunoblotting using a pSTAT3 or STAT3 antibody. Fold change of pSTAT3 over total STAT3. Data is representative of n = 3 - 4 independent experiments, respectively. Mean ± s.e.m. shown. *P* values were calculated using a repeated measures one-way ANOVA with a Tukey’s multiple comparisons test. * *P* ≤ 0.05, ** *P* ≤ 0.01, *** *P* ≤ 0.001. PonceauS or SimplyBlue SafeStains are shown for each representative membrane. HBVP = human brain vascular pericytes. SFM = serum-free media.

To determine whether ER+ breast cancer cells respond to IL-6 stimulation, we treated the two cell lines with the lowest basal STAT3 activation, MCF-7 and T-47D, with increasing concentrations of recombinant human IL-6 for 24 hours. STAT3 activation was detected in both MCF-7 and T-47D cell lines following IL-6 addition (Fig. 3b and 3c). In the MCF-7 cell line, we observed a positive trend in STAT3 activation for 2.5 ng/mL IL-6 treated cells compared with serum-starved control (*P* = 0.0980), with a statistically significant increase observed at 5.0 ng/mL (*P* = 0.0388) (Fig. 3b). Similarly, T-47D cells exhibited increased STAT3 phosphorylation at 2.5 ng/mL (*P* = 0.0087) and 5.0 ng/mL IL-6 (*P* = 0.0116) relative to control (Fig. 3c). Based on the findings, there was a positive trend indicating that STAT3 activation is IL-6 concentration-dependent (Fig. 3b and 3c; MCF-7: *P* = 0.1124 and T-47D: *P* = 0.7476). Overall, these data demonstrate that ER+ breast cancer cell lines with low basal STAT3 activity retain functional IL-6/JAK/STAT3 signaling and respond to exogenous recombinant human IL-6 protein stimulation.

### Docetaxel does not alter STAT3 activation in ER+ breast cancer cell lines

As breast cancer is frequently treated with docetaxel (DTX) at early- and advanced-stages, we assessed whether DTX alone activates STAT3 signaling in the selected ER+ breast cancer cell lines: MCF-7 and T-47D. DTX-treated ER+ breast cancer cells had minimal to no STAT3 activation after a 24-hour exposure compared to the positive control, 5 ng/mL IL-6 treatment group (Fig. 4a and 4b). A statistically significant increase in STAT3 activation was demonstrated between IL-6 against vehicle, and DTX-treated MCF-7 (Fig. 4a) and T-47D cells (Fig. 4b), respectively (MCF-7 vehicle: *P* = 0.001 and DTX: *P* = 0.001; T-47D vehicle: *P* = 0.0175 and DTX: *P* = 0.0175). Combinations of vehicle or DTX with IL-6 treatment had no statistical difference in STAT3 activation compared to IL-6 treatment alone for both ER+ breast cancer cell lines (MCF-7 vehicle: *P* = 0.9991 and DTX: *P* = 0.3587; T-47D vehicle: *P* = 0.7371 and DTX: *P* = 0.6905). Collectively, this data shows that while DTX does not directly activate STAT3, it does not prevent IL-6-mediated STAT signaling in ER+ breast cancer cell lines.

**Fig. 4:**
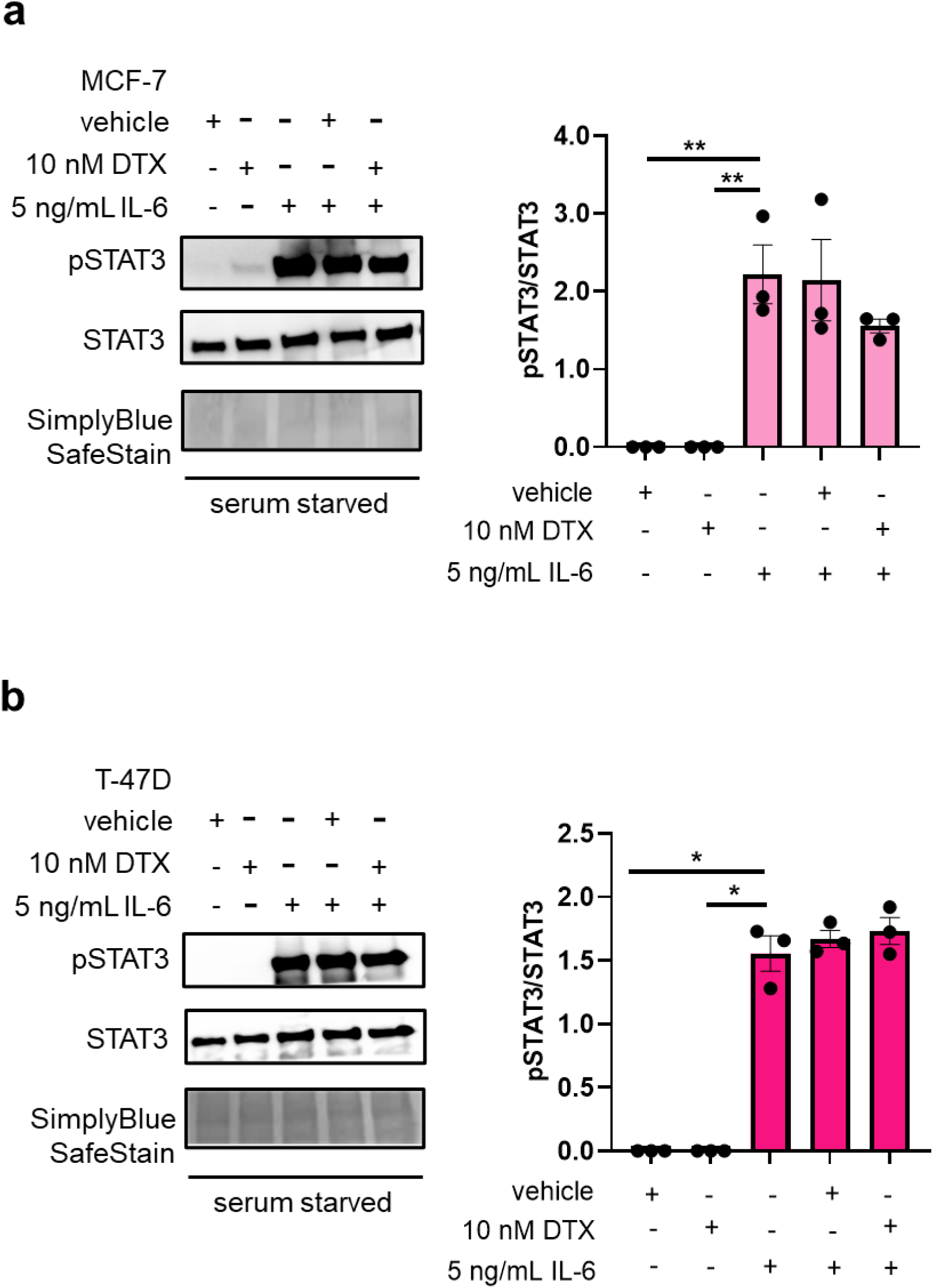
Docetaxel does not alter STAT3 activation in ER+ breast cancer cell lines. **a** MCF-7 and **b** T-47D cells were serum starved then incubated with vehicle, 10 nM DTX, or 5 ng/mL IL-6 alone or in combination as indicated for 24 hours. Cells were then lysed and subjected to immunoblotting using a pSTAT3 or STAT3 antibody. Fold change of pSTAT3 over total STAT3. Data is representative of n = 3 independent experiments. Mean ± s.e.m. shown. *P* values were calculated using a repeated measures one-way ANOVA with a Dunnett’s multiple comparisons test compared to the 5 ng/mL IL-6 group. * *P* ≤ 0.05, ** *P* ≤ 0.01. SimplyBlue SafeStains are shown for each representative membrane. DTX = docetaxel.

We next assessed whether DTX stimulates autocrine IL-6 production by ER⁺ breast cancer cells. DTX-treated MCF-7 cells had a significant change in IL-6 production at approximately 100 pg/mL following 24-hour treatment (Supplementary Fig. 3; *P* = 0.0025), a level more than ten-fold lower than that observed in DTX-treated HBVP cells (Fig. 1b). Vehicle-treated MCF-7 cells had IL-6 levels below the ELISA detection limit (3 pg/mL; *P* = 0.0025) compared to DTX-treated MCF-7 cells (Supplementary Fig. 3). In contrast, non-detectable concentrations were observed from T-47D cells treated with either vehicle or DTX (Supplementary Fig. 3; *P* > 0.9999). Together, the data demonstrate that DTX does not directly activate STAT3 in ER+ breast cancer cell lines and induces little to no autocrine IL-6 production.

### Tocilizumab reduces STAT3 activation induced by IL-6-enriched pericyte conditioned media in ER+ breast cancer cell lines

We examined the impact of DTX-stimulated HBVP cells on ER+ breast cancer cell lines by treating the MCF-7 and T-47D cell lines with conditioned media (CM) from DTX-treated HBVP cells, which is enriched in IL-6 (Fig. 1a and 1b), and evaluated STAT3 activation. When compared to serum-starved conditions, ER+ breast cancer cell lines treated with CM from DTX-treated HBVP cells had increased STAT3 activation (MCF-7: Fig. 5a, *P* = 0.0005, and T-47D: Fig. 5b, *P* ≤ 0.0001). STAT3 activation was similar among ER+ breast cancer cell lines treated with CM from DTX-treated HBVP cells compared to vehicle-treated HBVP cells (MCF-7: Fig. 5a, *P* = 0.4964, and T-47D: Fig. 5b, *P* > 0.9999), indicating that the basal level of IL-6 secretion from pericytes is sufficient to stimulate STAT3 signaling in ER+ breast cancer cells.

**Fig. 5:**
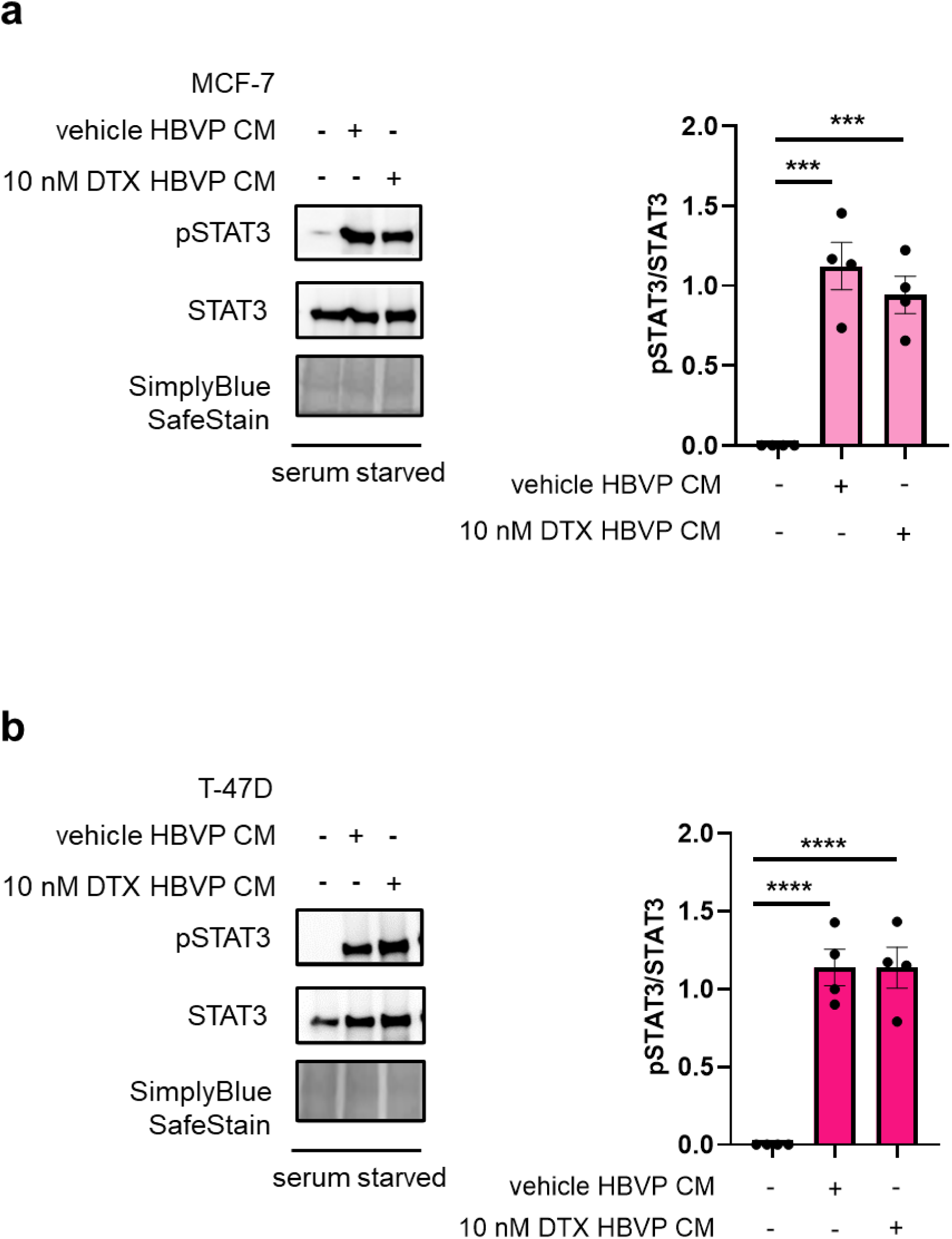
Conditioned media from docetaxel-treated pericytes induces STAT3 activation in ER+ breast cancer cells. **a** MCF-7 and **b** T-47D cells were serum starved then incubated with HBVP CM from vehicle or HBVP CM from 10 nM DTX as indicated for 24 hours. Cells were then lysed and subjected to immunoblotting using a pSTAT3 or STAT3 antibody. Fold change of pSTAT3 over total STAT3. Data is representative of n = 4 independent experiments. Mean ± s.e.m. shown. *P* values were calculated using an ordinary one-way ANOVA with a Tukey’s multiple comparisons test. *** *P* ≤ 0.001, **** *P* ≤ 0.0001. SimplyBlue SafeStains are shown for each representative membrane. HBVP = human brain vascular pericytes. CM = conditioned media. DTX = docetaxel.

Since CM from DTX-treated HBVP cells is enriched in IL-6 (Fig. 1a and 1b), we next tested whether STAT3 activation was mediated through IL-6 signaling. Tocilizumab is an FDA-approved humanized monoclonal antibody that inhibits IL-6 signaling by binding to soluble IL6-Rα (sIL-6Rα) and membrane-bound IL-6Rα (mIL-6Rα), resulting in blocking upstream of the IL-6/JAK/STAT3 pathway^70,77–80^. 0.1 µg/mL tocilizumab was chosen as the specific concentration was predicted to saturate the IL-6Rα binding site^78^. Tocilizumab significantly reduced STAT3 phosphorylation in both MCF-7 (*P* = 0.0068) and T-47D cell lines (*P* = 0.0334), compared with IgG-treated controls (Fig. 6a and 6b), indicating that CM-induced STAT3 activation is IL-6 dependent.

**Fig. 6:**
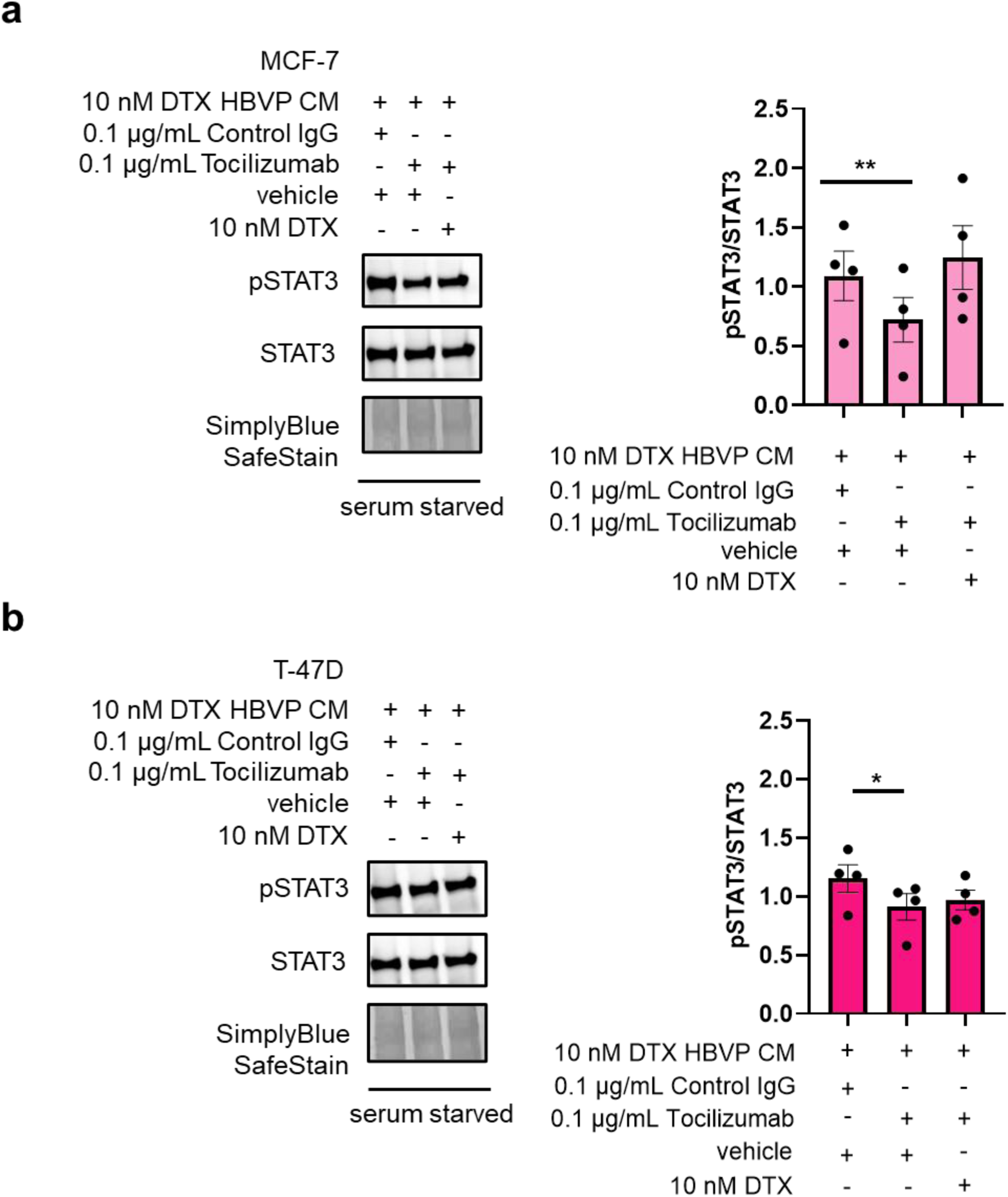
Tocilizumab reduces STAT3 activation induced by conditioned media from docetaxel-treated pericytes in ER+ breast cancer cell lines. **a** MCF-7 and **b** T-47D cells were serum starved then incubated HBVP CM from 10 nM DTX with 0.1 µg/mL control IgG, 0.1 µg/mL tocilizumab, vehicle, or 10 nM DTX in combination as indicated for 24 hours. Cells were then lysed and subjected to immunoblotting using a pSTAT3 or STAT3 antibody. Fold change of pSTAT3 over total STAT3. Data is representative of n = 4 independent experiments. Mean ± s.e.m. shown. *P* values were calculated using a repeated measures one-way ANOVA with a (**a**) Sidak’s and (**b**) Tukey’s multiple comparisons test. * *P* ≤ 0.05, ** *P* ≤ 0.01. SimplyBlue SafeStains are shown for each representative membrane. HBVP = human brain vascular pericytes, and CM = conditioned media. DTX = docetaxel.

We further assessed whether concurrent DTX treatment of tumor cells altered the effect of IL-6R blockade. Although a trend was observed, there was no statistically significant difference in STAT3 activation in the MCF-7 cell line treated with CM from DTX-treated HBVP cells, and tocilizumab with vehicle versus CM from DTX-treated HBVP cells, and tocilizumab with DTX (*P* = 0.1249) (Fig. 6a). In contrast, this trend was not observed in the T-47D cell line (Fig. 6b; *P* = 0.7362). In fact, STAT3 activation levels were similar between the T-47D cell line treated with CM from DTX-treated HBVP cells, and tocilizumab with vehicle versus CM from DTX-treated HBVP cells, and tocilizumab with DTX (Fig. 6b). The modest difference in MCF-7 cells may reflect low-level autocrine IL-6 secretion induced by DTX in this cell line (Supplementary Figure 3), which is absent in T-47D cells, potentially contributing to residual STAT3 activation despite IL-6R blockade.

Lastly, when comparing STAT3 activation in the MCF-7 cell line treated with CM from DTX-treated HBVP cells, and control IgG with vehicle against CM from DTX-treated HBVP cells, and tocilizumab with DTX, no statistically significant difference was observed (*P* = 0.7548), indicating similarity between the STAT3 activation levels (Fig. 6a). The T-47D cell line also had similar findings between the two different treatment groups (Fig. 6b; *P* = 0.0956). Together, these data suggest that tocilizumab is effective in decreasing STAT3 activation in ER+ breast cancer cell lines, even in the presence of DTX.

### In patient-derived ER+ breast cancer organoids, combined treatment with docetaxel and tocilizumab inhibits organoid growth, reduces STAT3 activation and cell proliferation, and increases apoptosis

High *IL6R* expression is associated with lower survival in treatment-naïve ER+ breast cancer patients, further supporting the idea that IL-6 signaling contributes to aggressive tumor behavior (Supplementary Fig. 4)^76,81^. To investigate this pathway in a translationally relevant model, we examined RNA-seq data from zero-passage patient-derived ER+ breast cancer organoids^82^ and found that this model largely retains *IL6R* expression (Supplementary Fig. 5a), whereas *IL6* expression is selectively preserved in a subset of cases in a non-exclusive way (Supplementary Fig. 5b). This retention of *IL6* and *IL6R* supports the use of these organoids as an *ex vivo* model to test IL-6-targeted therapies, including DTX and tocilizumab.

Combination treatment produced model-dependent effects. In Case 178, tocilizumab with DTX followed Bliss independence model predictions^83^ (Bliss excess = 0.000), showing no detectable synergy and minimal growth changes relative to DTX alone (Fig. 7a). In contrast, Case 176 exhibited a marked reduction in growth under combination treatment, resulting in a negative Bliss excess (–0.166) consistent with moderate synergy (Fig. 7b), while Case 179 showed a modest enhancement of growth suppression (Bliss excess = –0.050), indicative of mild synergy (Fig. 7c). Immunofluorescence staining for proliferation (EdU), IL-6 signaling (phosphorylated STAT3), apoptosis (NucView488; Caspase-3 activity reporter), mirrored these growth phenotypes. pSTAT3 signaling was low or undetectable in Case 178 (Fig. 7d), whereas tocilizumab-containing regimens completely suppressed endogenous pSTAT3 staining in Case 179 (Fig. 7e). Tocilizumab treatment did not affect Case 178 proliferation, but it decreased EdU positivity in Case 179. DTX treatment resulted in apoptosis activation in both cases. Together, these data indicate that tocilizumab enhances DTX efficacy in a subset of patient-derived models, with its combinatorial benefit correlating with STAT3 signaling suppression, highlighting *IL6R* high ER+ breast cancers as a potential target population for IL-6-directed combination therapy.

**Fig. 7:**
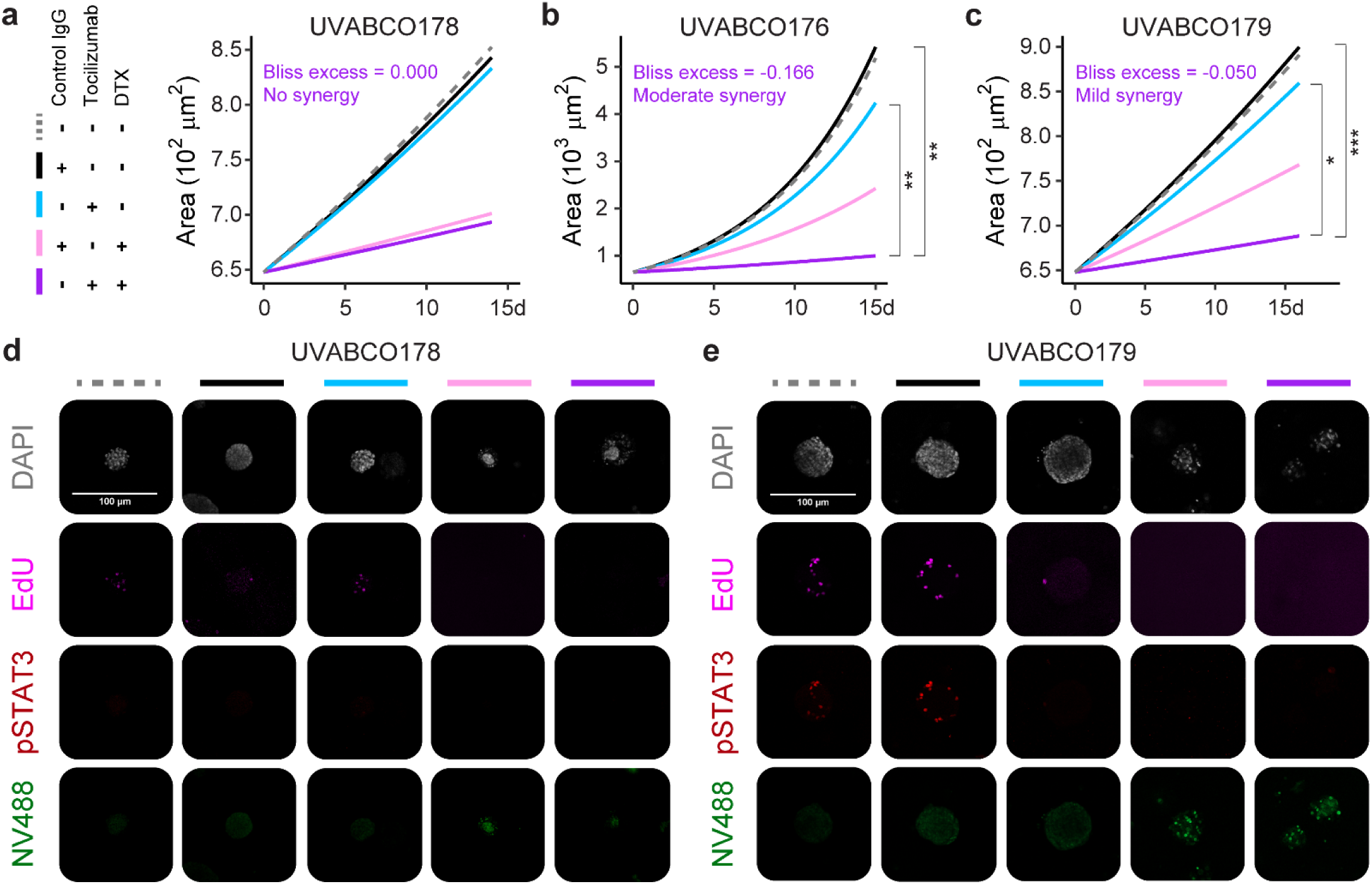
Model-dependent synergy between tocilizumab and docetaxel in zero-passage patient-derived ER+ breast cancer organoids. (**a**–**c**) Longitudinal spheroid growth of UVABCO178 (**a**), UVABCO176 (**b**), and UVABCO179 (**c**) over 14-16 days following treatment with control IgG (black), tocilizumab (0.1 µg/mL; cyan), DTX (10 nM; light pink), or the tocilizumab + DTX combination (purple). Lines represent model-fitted exponential growth curves for spheroid area over time, derived from linear regression of log-transformed area (log(Area) ∼ day) (n = 8 – 780 organoids per patient, time point, and condition). Bliss synergy excess for the tocilizumab + DTX combination are reported in each panel. Statistical significance between treatment groups is indicated (* *P* ≤ 0.05, ** *P* ≤ 0.01, *** *P* ≤ 0.001).(**d**–**e**) Representative immunofluorescence images of spheroids from UVABCO178 (**d**) and UVABCO179 (**e**) following treatment with control IgG, tocilizumab, DTX, or the combination. Staining includes DAPI (nuclei, white), EdU (proliferation, magenta), pSTAT3 (IL-6 signaling, red), and NucView488 (NV488; apoptosis, green). Scale bars, 100 µm. DTX = docetaxel.

## Discussion

Microtubule targeting agents (MTAs) have been the standard of care but are often underappreciated for their contribution to therapeutic efficacy in combination with other drug treatments, as demonstrated by multiple cancer clinical studies^84,85^. Docetaxel (DTX), an MTA, is a mainstay in the treatment of adult solid tumors, including breast cancer, and is initially highly effective. Unfortunately, patients with ER+ breast cancer have a high likelihood of recurrence, with one explanation being the unappreciated role of the TME and the presence of inflammatory cytokines^86–88^. The effects of DTX on cancer cells have been well studied; however, the effects of DTX directly on pericyte-mediated inflammation are not well-defined in mediating breast cancer chemoresistance. To address this question, our study focused on using DTX-treated pericytes to 1) identify what cytokines are enriched in their secretome, and 2) elucidate the role of the pericyte-derived secreted factors in therapeutic resistance. We hypothesized that the pericyte secretome consisted of inflammatory cytokines that would contribute to cancer cell chemoresistance.

Here, we show that IL-6 production from pericytes is induced with a concentration as low as 10 nM DTX after 24-hour treatment *in vitro* (Fig. 1). 10 nM DTX is a low, biologically active concentration in experimental models, distinct from the much higher doses used in patients. As such, this approach may encourage chemoresistance as these concentrations may also affect secreted products within the TME. In more recent studies, low doses of therapeutics have been utilized to reduce off-target effects and decrease the likelihood of generating senescent, resistant, and drug-tolerant tumor persister cells^89–92^. We then validated that our breast cancer cell lines had the IL-6R complex subunits at the cell surface, ensuring that IL-6 secreted by pericytes could effectively signal in these cells (Fig. 2). Using two ER+ breast cancer cell lines, MCF-7 and T-47D, with low constitutive STAT3 activation, we proved that exogenous IL-6 upregulates STAT3 signaling (Fig. 3). DTX alone does not appear to alter STAT3 activation in the MCF-7 and T-47D cell lines (Fig. 4). When adding CM from DTX-treated pericytes onto cancer cells, we observed robust STAT3 activation (Fig. 5). To further probe the role of IL-6 in this effect, we introduced tocilizumab (IL-6R inhibitor) at 0.1 µg/mL, which resulted in a reduction of STAT3 phosphorylation. Collectively, we did not see a complete reduction with tocilizumab (Fig. 6), potentially due to the presence of different secreted factors that can activate the JAK/STAT3 signaling pathway, such as IL-6 family cytokines, IL-10 family cytokines, growth factors/hormones, and other potential activators in the pericyte CM^93,94^. Moreover, we selected a low clinical concentration for tocilizumab, which could also explain the incomplete reduction in STAT3 activation^95,96^. The combination of docetaxel and tocilizumab acted synergistically in zero-passage patient-derived ER+ breast cancer organoids that exhibited autocrine IL-6 signaling, resulting in decreased proliferation, decreased pSTAT3, and increased apoptosis, compared to untreated, control conditions, and single treatments alone (Fig. 7).

We also showed that cells from Supplementary Table 1 did not have similar levels for *IL6R* mRNA and at the cellular surface for IL-6Rα compared to the T-47D cell line, which had high levels of *IL6ST* mRNA and IL-6Rβ for cell surface abundance, accordingly (Fig. 2). This finding could be explained by IL-6Rβ being ubiquitously expressed^97^ and how differential shedding occurs for the two different subunits, where IL-6Rβ sheds at almost negligible levels in comparison to IL-6Rα^98^. Furthermore, *IL6R* expression could result in both membrane-bound IL-6Rα (mIL-6Rα) and soluble IL-6Rα (sIL-6Rα). sIL-Rα can be generated via alternative splicing, in addition to a more recently discovered form of the sIL-6Rα with a novel C-terminus generated by proteolytic cleavage (N-glycosylation); therefore, a potential reason to explain the discrepancy between mRNA levels and the abundance at the cellular surface for the receptor subunits^99^. Non-tumorigenic breast epithelial cell lines (MCF-10A and 184A1) had relatively lower levels of the IL-6Rβ subunit at the cell surface relative to the HBVP cells, HCC1500, and T-47D cell lines, but had similar IL-6Rα cellular surface abundance (Fig. 2b). This observation further provides evidence that other stromal cell types in the TME, even non-tumorigenic breast epithelial cells, are capable of taking up IL-6 from the TME from other cell types through paracrine IL-6 signaling and promoting cancer progression through epithelial-mesenchymal transition and stemness^100^.

Intriguingly in the MCF-7 cell line, the combination of tocilizumab and DTX with DTX-treated pericyte CM had similar levels of STAT3 activation compared to DTX-treated pericyte CM with control IgG and vehicle (Fig. 6a). On the contrary, this observation was not noted for the T-47D cell line (Fig. 6b). We speculate that the trend was observed, in part, because perhaps DTX induces IL-6 production in the MCF-7 cell line to a much greater extent than the T-47D cell line (Supplementary Fig. 3). This suggests that there could be feedback (autocrine signaling) of exogenous IL-6 that recovers the pSTAT3 reduction with just tocilizumab treatment alone.

Our findings show that combining tocilizumab with DTX improved therapeutic efficacy in zero-passage patient-derived ER+ breast cancer organoids with intact IL-6/STAT3 signaling (Fig. 7). The mild to moderate synergy observed *ex vivo* suggests that both autocrine, tumor-intrinsic, and paracrine, TME-specific IL-6 signaling may contribute to DTX sensitivity. Given that tocilizumab-sensitive organoids from two independent patients exhibited negative Bliss excess values, the posterior belief that the interaction is synergistic is increased relative to a neutral prior. Our results extend previous studies by showing that targeting the IL-6 pathway can sensitize clinically relevant, patient-derived *ex vivo* models to chemotherapy, underscoring its translational potential for therapeutic regimens in ER+ breast cancer (Fig. 7).

In conclusion, our work demonstrates that DTX treatment directly stimulates pericytes in the TME to enhance IL-6 production (Fig. 8). This finding highlights that therapeutics could have off-target effects depending on drug concentration and treatment duration. This study further suggests that the combination of DTX with an IL-6R inhibitor may be an effective treatment for ER+ breast cancer patients to reduce chemoresistance and recurrence, as shown by the reduction of STAT3 activation (Fig. 6, 7, and 8). Previous work utilized tocilizumab as an inhibitor for IL-6R in TNBC models, without the inclusion of pericytes, and without ER+ breast cancer models^101^. Future work could utilize omics approaches to discover novel secreted factors derived from pericytes with and without therapeutic exposure, to gain a deeper understanding of how the pericyte secretome contributes to cancer progression, therapeutic resistance, and metastasis. The use of tocilizumab in our study as an additional FDA-approved therapeutic with the standard of care, added sequentially and/or in combination, suggests and emphasizes that breast cancer treatment may require multiple targets to overcome recurrence and chemoresistance. Furthermore, administering lower doses of different therapeutics in patients could work synergistically in anti-tumor efficacy and be more effective in overcoming resistance^102,103^. In summary, elucidating the mechanisms behind how stromal cell types like pericytes contribute to drug resistance might lead to novel therapeutic approaches to overcome resistance for breast cancer patients in the clinic.

**Fig. 8:**
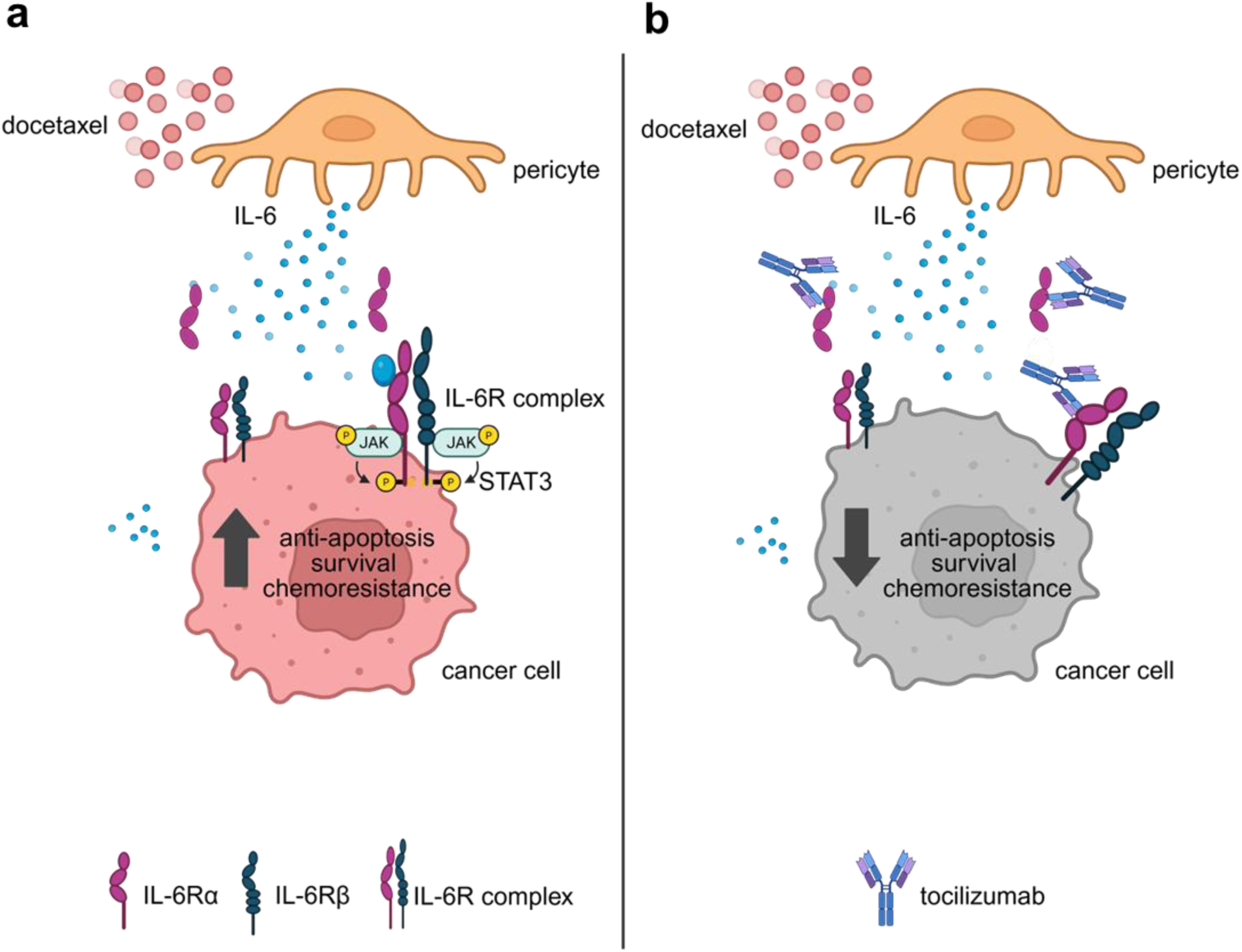
Proposed model of docetaxel-induced pericyte signaling and IL-6R targeted intervention in ER+ breast cancer. **a** Docetaxel directly impacts pericytes within the tumor microenvironment, inducing secretion of IL-6. Pericyte-derived IL-6 engages the IL-6R complex (IL-6Rα/IL-6Rβ) on ER+ breast cancer cells, activating JAK/STAT3 signaling and promoting anti-apoptotic signaling, cancer cell survival, and chemoresistance. **b** Pharmacologic inhibition of IL-6R with tocilizumab disrupts IL-6 mediated signaling between pericytes and cancer cells. IL-6R blockade attenuates STAT3 activation, resulting in reduced anti-apoptotic signaling and decreased chemoresistance in ER+ breast cancer cells treated with docetaxel. Generated from BioRender.

### Limitations and strengths of the study

Although these results present rigorous and reproducible findings, there are limitations to this study: (1) We utilized an *in vitro* model of pericytes (human brain vascular pericytes), in which the pericytes are not derived from the breast, (2) Conditioned media (CM) from pericytes (indirect co-culture) instead of a direct co-culture with both pericytes and cancer cell lines was used. Although we just examined the CM of pericytes, pericytes are similar to other cell types, such as adipocytes, fibroblasts, and immune cells, whose secretome includes IL-6 production. However, the amount of IL-6 production would be dependent on certain variables and conditions. Biologically, we recognize that there are more secreted factors than IL-6 from pericytes that could activate the JAK/STAT3 signaling pathway in ER+ breast cancer cells. Tocilizumab has also been shown to inhibit or reduce other cytokines (IL-8) at certain concentrations and specific time points, as well as impact angiogenesis^104^. (3) DTX was utilized as the sole chemotherapeutic as a microtubule targeting agent and microtubule stabilizer. MTAs consist of microtubule stabilizers and destabilizers, and it has been shown that MTAs are not all the same in their mechanisms of action and have unique anti-inflammatory and pro-inflammatory effects on different cell types in the TME^60,105–107^. DTX gets detected by cells as an inflammatory trigger through pattern recognition receptors like the Toll-like receptor 4 (TLR4), which results in IL-6, granulocyte-colony stimulating factor (G-CSF)^108^, and tumor necrosis factor alpha (TNFα) secretion^109^ as a few examples, and ultimately leads to therapy resistance^110–112^. Further, there is evidence that DTX upregulates Toll-like receptor 4 (TLR4) mRNA expression, but not activation of the TLR4/NF-κB (nuclear factor kappa-light-chain-enhancer of activated B cells) pathway, which promotes the expression of pro-inflammatory cytokines^113^. Paclitaxel, the first taxane discovered and another MTA, binds to the TLR4-accessory molecule, MD-2, and activates the TLR4 pathway in murine cells^114^. Molecular docking studies conducted found that paclitaxel binds to MD-2 in human cells, and its binding site is similar to lipopolysaccharide (LPS), the primary ligand of TLR4, resulting in decreased LPS signaling^115,116^. Thus, it would be intriguing and of value to assess the differential effects of a panel of MTAs on the pericyte secretome in combination with an IL-6R inhibitor on anti-tumor efficacy.

Our study has several notable strengths: (1) We utilized primary human pericytes that are of woman origin rather than murine-derived model, which enhances the physiological relevance of our findings. (2) The use of brain-derived pericytes may provide additional relevance to metastatic ER+ breast cancer, especially considering the propensity of this subtype to metastasize to the brain. (3) We employed physiologically relevant drug concentrations, including clinically informed low doses of docetaxel and tocilizumab, aligning our *in vitro* and *ex vivo* models more closely with therapeutic exposure scenarios. (4) In addition to using pericyte CM, we performed parallel experiments with recombinant human IL-6 alone to isolate the contribution of a singular cytokine and strengthen mechanistic specificity. (5) We leveraged miniaturized zero-passage patient-derived ER+ breast cancer organoids, which preserve estrogen receptor expression and enable the testing of multiple drug combinations within material derived from a single patient^82^. (6) Our study is also distinct from recent studies in the field. He et al.^117^ studied pericyte phenotype switching, whereas our study interrogates how therapy-altered pericytes influence tumor cell signaling and therapeutic response. (7) Lastly, our work utilizes CM acquired from DTX-treated pericytes, which looks at the effects of pericyte crosstalk onto cancer cells, instead of just evaluating the effects of eribulin, a microtubule destabilizer, and tocilizumab in a drug-resistant model, as conducted by Hattori et al.^118^ (2025). By focusing on treatment-induced pericyte–tumor crosstalk, our study provides a TME-centered perspective on mechanisms of chemoresistance.

## Methods

### Reagents and Materials

Tocilizumab (#A2012) and human IgG isotype control (#A2051, RRID:AB_3096062)) were purchased from SelleckChem, while docetaxel (DTX) (#01885-5MG-F) and bovine serum albumin (#A193325G) were purchased from Sigma Aldrich. DMSO, which is the vehicle control solvent for DTX was purchased from ThermoFisher (#J66650.AK). In addition, HyClone™ Water, Molecular Biology Grade (#SH3053802) was bought from Cytiva. Milk powder (#NC9952266) and Ponceau S staining solution (#A40000279) were purchased from ThermoFisher for immunoblotting. Recombinant human IL-6 protein was purchased from Peprotech (#200-06-5UG or 20UG).

The following antibodies were purchased from Cell Signaling Technology for immunoblotting: Phospho-Stat3 (Tyr705) (D3A7) XP® (#9145, rabbit mAb, RRID:AB_2491009), STAT3 (124H6) (#9139, mouse mAb, RRID:AB_331757), and secondary anti-mouse IgG, HRP-linked antibody (#7076, RRID:AB_330924), and secondary anti-rabbit IgG, HRP-linked antibody (#7074, RRID:AB_2099233).

Antibodies used for flow cytometry studies were purchased from both BioLegend and BD Biosciences. For IL-6Rα and IL-6Rβ cell surface detection, we purchased APC anti-human CD126 (IL-6Rα) (BioLegend #352806, RRID:AB_11204258) and PE anti-human CD130 (gp130) (BioLegend #362004, RRID:AB_2563402).

### Cell Culture

T-47D (HTB-133, RRID:CVCL_0553), MCF-7 (HTB-22, RRID:CVCL_0031), and HCC1500 (CRL-2329, RRID:CVCL_1254) ER+ breast cancer cell lines were obtained from ATCC. The MDA-MB-231 (HTB-26, RRID:CVCL_0062) cell line was obtained from Dr. Jinyang Li in Dr. Anthony Kossiakoff’s laboratory (University of Chicago). 184A1 (CRL-8798, RRID:CVCL_3040) and MCF-10A (CRL-10317, RRID:CVCL 0598) mammary epithelial cell lines were provided by Dr. Marsha Rosner (University of Chicago). Human Brain Vascular Pericytes (HBVP), also labelled as pericytes throughout the manuscript, were purchased from ScienCell Research Laboratories (#1200; lot number 32562, previously #1020). All fetal bovine serum (FBS) used for cell culture was heat-inactivated (Life Technologies #A5669701; Thermo Fisher Scientific #1600-044), except for pericyte medium.

T-47D and MDA-MB-231 cell lines were cultured in Dulbecco’s Modified Eagle Medium (DMEM) (Thermo Fisher Scientific #10-013-CV) supplemented with 10% FBS and 1% penicillin-streptomycin (P/S; Gibco #15-140-122). Initially, the MDA-MB-231 cell line was cultured in 1:1 DMEM supplemented with 10% FBS and 1% P/S and Gibco™ Roswell Park Memorial Institute (RPMI) 1640 medium (Thermo Fisher Scientific #11-875-119) with 10% FBS and 1% P/S. The MCF-7 cell line was initially maintained in Improved Minimum Essential Medium (IMEM) (Corning #A1048901) with 10% FBS and 2.5 mL of 10 mg/mL gentamicin (Gibco #15710064) in 500 mL, and subsequently grown in DMEM supplemented with 10% FBS and 1% P/S. 184A1 and MCF-10A cell lines were cultured following manufacturer recommendations in MEGM™ Mammary Epithelial Cell Growth Medium BulletKit™ (Lonza #CC-3150). HBVP cells were maintained in ScienCell Pericyte Medium (ScienCell Research Laboratories #1201) and were passaged up to a maximum of passage 10 based on morphology and overall cell health. The HCC1500 cell line was cultured in Gibco™ RPMI 1640 medium with 10% FBS and 1% P/S. All cancer cell lines were cultured at 37°C in a humidified incubator with 5% CO₂ and were passaged up to passage 40 before disposal. All lines were routinely tested for mycoplasma contamination (Invivogen #rep-mys-50).

### Cell Counting Kit-8 assay

Approximately 1000 MCF-7 cells/well, 4000 MDA-MB-231 cells/well, 7000 T-47D cells/well, and 5000 HBVP cells/well were seeded into a 96-well plate using their complete media, respectively. Cells were allowed to adhere overnight. For Supplementary Figure 1, cells were treated with 10 nM DTX for 24 hours at 37°C, 5% CO_2_ in a total volume of 100 µL in technical triplicates for n = 3 independent experiments. Following 24 hours, 10 µL CCK-8 (Dojindo #CK04-11; Thermo Fisher Scientific #NC9864731) was added to each well for 3 to 4 hours. The 96-well plates were read at 450 nm using the SpectraMax i3x platereader (Molecular Devices). % control was calculated using the following formula: (experimental sample absorbance value at 450 nm minus media background) divided by vehicle-treated cells absorbance at 450 nm minus media background), followed by multiplying by 100.

### Zero-passage patient-derived ER+ breast cancer organoids

#### Initiation and culture

ER+ breast cancer patient-derived organoids were established and cultured as previously described^82^. Shortly, three 50-mL conical tubes, each containing two glass slides of tumor scrapes, were centrifuged at 450 rcf for 5 minutes at 8°C. The pellet was washed with D-BSA (DMEM GlutaMAX (Gibco #10569010)) with 0.1% fatty acid-free BSA (Sigma #A6003) and transferred to a 1.5 mL tube. Upon three subsequent washes, the pellet was digested for 15 minutes at 37°C with shaking (350 rpm) in Type 1 medium containing collagenase II (Fisher #17101015; final concentration = 1 mg/mL) and ROCK inhibitor (StemCell #72304; final concentration = 10 μM). Digestion was stopped by adding FBS (Gibco #16170086; 1/10th volume), followed by vigorous pipetting with a 1 mL micropipette to fragment the tissue. If required, the mixture was strained through a pre-wet 100 μm strainer (VWR #76327102), washed twice with D-BSA, and incubated with RBC lysis buffer (Sigma #11814389001) for 2 minutes at room temperature (RT) before the final wash. The resulting pellet was suspended in Matrigel (Corning #354230) at a volume ratio of 4:1 Matrigel:cell pellet and pipetted as single 5 μL drops per well of a 96-well plate. The plate was incubated upside down for 20 minutes at 37°C before adding 100 μL of pre-warmed Type 2 medium^119^ to the solidified drops in each well. Organoid cultures were kept at 37°C in a humidified incubator with 5% CO_2_, and the medium was exchanged every 2–5 days by gentle aspiration and pipetting.

#### Drug treatment

Medium was removed at day 5 and replaced with media containing DMSO (final concentration = 0.1%) or 10 nM DTX, human IgG isotype control, or tocilizumab (0.1 μg/mL). The fresh medium ± drug was added on day 7, exchanged on day 10, and added on day 12.

#### Organoid growth measurement

Organoid images were digitally acquired every 2–3 days with a 2×/0.08 NA long working-distance plan apochromat objective on an EVOS M7000 (ThermoFisher #AMF7000) with DiamondScope software and analyzed with OrganoSeg2^120^. The image sets were segmented using the following parameters: out-of-focus correction, split whole image, edge correction, intensity threshold = 1.5(176)/0.8(178)/1.25(179), size threshold = 100, window size = 100, contaminant intensity=0.25(176)/0.15(178&179), minimum circularity=0.4(176)/0.15(178&179). Area measurements were exported as an .xls file and further analyzed in R Studio.

#### EdU treatment and pSTAT3 immunostaining

Organoid proliferation was assessed using a 5-ethynyl-2′-deoxyuridine (EdU) incorporation assay to measure DNA synthesis. EdU (Invitrogen #E10187; final concentration 50 μg/mL) and NucView488 (Sigma #SCT100; final concentration = 2.5 μM) were added at the endpoint of culture and incubated overnight at 37°C. Organoids were then fixed in 3.7% paraformaldehyde overnight at 4°C, permeabilized with 0.1% Tween in PBS for 1 hour at RT, and processed for EdU detection using Click-it! Reaction. 100 μL of 1 M Tris-HCl pH 8.5, 1 M CuSO4, Alexa Fluor 647 azide triethylammonium salt (Invitrogen #A10277; final concentration 2 μg/mL), and 0.5 M ascorbic acid (Sigma #A7631) was added to each well and incubated for 1 hour at RT. Then, organoids were blocked for 1 hour at RT in blocking solution (1x Western Blocking reagent (Roche #11921673001), diluted in PBS, and supplemented with 0.3% Triton X-100. Primary p-STAT3 (Tyr705) (Cell Signaling Technology #9145, RRID:AB_2491009) antibody diluted 1:200 in blocking solution was added overnight at 4°C. The next day, wells were washed with PBS three times for 5 minutes, and Alexa Fluor 555 secondary antibody (Thermo Fisher Scientific #A-21428, RRID:AB_2535849) diluted 1:200 in blocking solution was added for 1 hour at RT. DAPI (Invitrogen, #D1306; final concentration = 0.5 μg/mL) was added to each well for 5 minutes and washed twice with PBS, after which the final 100 μL of PBS was added. Images were acquired on a Zeiss confocal system (LSM900) and processed using Fiji software (ImageJ version 2.16.0).

### Conditioned media preparation and storage

#### For use in immunoblotting experiments

When cells hit a confluency of 70-90%, cells were washed with PBS prior to the addition of serum-free media (0.1% Bovine serum albumin (Millipore Sigma/Roche #3117057001), 1% P/S (Gibco #5-140-122)) for 24 hours. Conditioned media were transferred to a 15 mL or 50 mL conical tube, or a 1.5 mL Eppendorf tube, and centrifuged for 5 minutes at 600 g at RT. The supernatant was transferred to a new conical tube or Eppendorf tube and stored at -80°C or utilized immediately.

For cells treated with vehicle or 10 nM DTX for 24 hours in their respective complete media, they were treated at a confluency of 70-80%, followed by PBS washes, and then addition of serum-free media (0.1% BSA (Millipore Sigma/Roche #3117057001), 1% P/S for 24 hours prior to collection. Thus, the conditioned media were collected after 48 hours from drug exposure. This experimental protocol was conducted due to collecting conditioned media without drug presence and drug degradation to add onto the cancer cells^121^. Conditioned media were transferred to a conical tube or Eppendorf tube and centrifuged for 5 minutes at 600 g at RT. The supernatant was transferred to a new conical tube or Eppendorf tube and stored at -80°C or utilized immediately. All cells used DMEM serum-free media (SFM), which consists of DMEM, 0.1% BSA, and 1% P/S, except for the HCC1500 cells, which used RPMI SFM (RPMI, 0.1% BSA, and 1% P/S).

6-well plates were utilized for cells, where up to 1 mL conditioned media was harvested and approximately 600 µL frozen down at -80°C. T-75 and T-175 flasks were also used in which 8 mL conditioned media was harvested, and 24-25 mL conditioned media was harvested, respectively.

#### For cytokine array and ELISA conditioned media preparation

6-well plates were used where 1 mL SFM was added and about 600 µL of conditioned media was frozen down at -80°C.

When cells hit a confluency of 70-80%, cells were treated with vehicle or 10 nM DTX for 24 hours in their complete media, followed by PBS washes, and then addition of SFM with 1% P/S for 24 hours. Conditioned media were then transferred to a conical tube or Eppendorf tube and centrifuged (600 g, 5 minutes, RT). Without disturbing the pellet, the supernatant was transferred to a new conical tube or Eppendorf tube and stored at -80°C or utilized immediately for experiments.

### Flow cytometry

All flow cytometry was performed at the Cytometry and Antibody Technology Facility at University of Chicago, which receives financial support from the Cancer Center Support Grant (P30CA014599). RRID: SCR_017760. For cell surface detection only: Cells were grown until confluence reached up to 90%. For each sample, cells were harvested using 0.25% trypsin-EDTA, phenol red (Thermo Fisher Scientific #25-200-056) or 0.05% trypsin-EDTA (Thermo Fisher Scientific #25-300-054), which was specifically used for the HBVP cells, and MCF-10A and 184A1 cell lines, accordingly. Media with serum was used to neutralize the 0.25% or 0.05% trypsin-EDTA, respectively, and then resuspended in flow buffer (Bio-Techne #FC001). Cells were spun down at 1000 rpm for 3 to 4 minutes and resuspended in 100 µL volume flow buffer in a 96-well round-bottom plate (Thermo Fisher Scientific #07-200-87). Following counting the number of cells with a countess (Countess 3; Thermo Fisher Scientific), approximately 0.5 million cells were stained with 5 µL of Human TruStain Fcx block (BioLegend #422302, RRID: AB_2818986) at RT in the dark for 10 minutes, followed by staining with 5 µL of either APC anti-human CD126 (IL-6Rα) (BioLegend #352806, RRID:AB_11204258) or PE anti-human CD130 (gp130) (BioLegend #362004, RRID:AB_2563402) for 20 minutes on ice in the dark. Cells were spun down at 400 g for 3 minutes at 4℃ at 100 µL volume in a 96-well round-bottom plate. The supernatant was aspirated, and the cell pellet was resuspended in at least 200 µL of flow buffer twice. Cells were then resuspended in 200 µL of flow buffer to run for flow cytometry analysis using the NovoCyte Penteon to examine PE and APC detection. Data was analyzed by FlowJo software (TreeStar Inc, RRID:SCR_008520). Two to three technical replicates were conducted for each cell line with both APC anti-human CD126 (IL-6Rα) and PE anti-human CD130 (gp130) (IL-6Rβ) for a total of n = 3 to 5 independent experiments. All samples were acquired in the University of Chicago Antibody and Cytometry Core Facility.

### Cytokine array

Cells were grown to 70-80% confluency and treated with vehicle or 10 nM DTX for 24 hours. Cells were then washed with PBS. The addition of 1 mL DMEM SFM or the cells’ respective media (HBVP cells) was conducted for 24 hours, and then the conditioned media were harvested. The conditioned media were collected after centrifuging for 5 minutes at RT and 600 g. The relative concentrations of 105 cytokines were measured using the Proteome Profiler Human XL Cytokine Array Kit (R&D Systems, part of Bio-Techne #ARY022B). Conditioned media were either used fresh, right away, or stored in 600 µL aliquots in the -80°C freezer. Multiple rounds of freeze-thaw were avoided. The manufacturer’s protocol was followed. Imaging was completed using the Bio-Rad Chemi Doc Imager, and quantification of average adjusted volumes was performed to generate fold change between conditioned media from vehicle and 10 nM DTX-treated pericytes using the Bio-Rad Chemi Doc software.

### ELISA

To analyze basal IL-6 secretion from untreated cells, cells were seeded in 6-well plates. Once cells hit a confluency of 80-90%, cells were washed with PBS, followed by incubation with 1 mL DMEM SFM or RPMI SFM, respectively, for 24 hours.

To assess the effects of docetaxel on IL-6 secretion, cells in 6-well plates were treated with vehicle or 10 nM DTX for 24 hours, followed by PBS washes, and incubation with 1 mL DMEM SFM for 24 hours.

To harvest conditioned media, the supernatant was transferred into 1.5 mL Eppendorf tubes or 15 mL conical tubes and centrifuged for 5 minutes at 2300 rpm, which is similar to 600 g at RT. Without disturbing the pellet at the bottom, the supernatant was then transferred to 1.5 mL Eppendorf tubes to be frozen down at -80°C or utilized fresh/immediately for running the ELISA using the Human IL-6 ELISA kit – Quantikine (Bio-Techne #D6050B) as per the manufacturer’s instructions. No freeze-thaw cycles occurred previously for these samples. The manufacturer’s protocol was followed with the following modification: for HBVP samples (vehicle or 10 nM DTX): Samples were run in technical duplicates with a minimum of n = 2 independent experiments. 450 nm absorbance values minus 540 nm absorbance values were utilized for wavelength correction using the SpectraMax i3x platereader (Molecular Devices).

### RNA isolation to RT-qPCR

#### RNA isolation and quantification

Total RNA from 1) pericytes treated with vehicle or 10 nM DTX for 24 hours and 2) a panel of cells and cell lines (Supplementary Table 1), respectively, was isolated using the RNeasy Mini Kit (Qiagen #74106), according to the manufacturer’s protocol for total RNA isolation. All preparation and handling of RNA took place under RNase-free conditions. Quality and quantity of RNA samples were assessed using a NanoDrop spectrophotometer (Thermo Fisher Scientific, RRID:SCR_015804).

#### cDNA synthesis

High-quality RNA samples were aliquoted (1 µg of total RNA) and used for the reverse transcriptase reaction using the Verso cDNA Synthesis Kit (Thermo Fisher Scientific #AB1453B) in the SimpliAmp Thermal Cycler (Thermo Fisher Scientific). For all reactions, the same amount of total RNA (1 µg/reaction) from the same isolation was used. All experiments (RNA isolation, cDNA synthesis, and real-time polymerase chain reaction (RT-qPCR)) were performed in technical duplicates, and a minimum of n = 3 independent experiments.

#### RT-qPCR reaction

RT-qPCR was performed using the TaqMan Fast Advanced Master Mix Protocol (Applied Biosystems by Life Technologies). The Taqman Fast Advanced Master Mix (Thermo Fisher Scientific #4444557) was mixed with diluted cDNA sample (5 ng total per sample), RNAse-free water (Cytiva #SH3053802), and primers. Primers used were Taqman Gene Expression Assays *IL6* (Hs00985638_m1, Life Technologies), *IL6ST* a.k.a. IL-6Rβ (Hs00174360_m1, Life Technologies), *IL6R* a.k.a. IL-6Rα (Hs01075666_m1, Life Technologies), and GAPDH (Hs02786624_g1, reference gene, Life Technologies). *GAPDH* was utilized as the reference gene. Samples were loaded onto a 200 µL 96-well PCR plate (MidSci #PR-PCR2196F), closed with the MicroAmp Optical Adhesive Film (Thermo Fisher Scientific #4311971), and run using the QuantStudio 3 Real-Time PCR System (Thermo Fisher Scientific). The RT-qPCR reaction was carried out for 40 cycles with UNG incubation at 50°C for 2 minutes, polymerase action at 95°C for two minutes, denaturation at 95°C (1 second), and annealing/extending at 60°C (20 seconds). For *IL6R* and *IL6ST* mRNA, samples were normalized to MDA-MB-231 samples at 1, while *IL6* mRNA samples were normalized to vehicle-treated HBVP cells at 1.

### Immunoblotting for basal STAT3 activation for untreated cells and STAT3 activation following stimulation

All cells (HBVP, MDA-MB-231, MCF-7, T-47D, and HCC1500) were cultured and seeded in 6-well plates using their respective complete media. Once 80-90% confluence was reached in 6 well plates, cells were washed 1x with ice-cold PBS. PBS was aspirated, followed by the addition of RIPA buffer (Sigma Aldrich #R0278) with inhibitors. Cells were incubated with RIPA buffer for 20 minutes on ice in the 6-well plates, followed by scraping with the back of a P1000 tip and/or cell scraper, followed by transferring lysate to pre-chilled and pre-labelled 1.5 mL Eppendorf tubes. Samples were vortexed 3 x 10 minutes on ice and spun down at 14000 g for 10 minutes at 4°C. The supernatant was transferred to new pre-chilled 1.5 mL Eppendorf tubes. BCA assays were conducted to generate normalized cell lysates amongst samples. Approximately 40 – 50 µg protein for each sample was loaded with 4x LDS buffer (Bio-Rad #1610747) and 2-Mercaptoethanol (Sigma Aldrich #M7154) into 4-20% Bio-Rad Mini-Protean TGX stain-free gels (10-well comb, 30-50 µL #4568094) with 7-8 µL Precision Plus Protein Dual Color Standards Ladder (Bio-Rad #1610374). The samples were run in the gel at 100V-120V and were then transferred for 1.5 hours at 90V on ice with ice-cold transfer buffer using PVDF membrane (Cytiva Amersham™ Hybond™ PVDF Membranes: Rolls; Thermo Fisher Scientific #45-004-021). Following transfer, the membranes were allowed to dry and then reactivated with methanol prior to staining with Ponceau S for 5 minutes at RT while shaking. After a quick DI water and 1x TBS wash, blocking with 5% BSA-TBST (for pSTAT3) or 5% Milk-TBST (STAT3) for 1 hour at RT, shaking was performed. For pSTAT3 detection, 1:1000 of p-STAT3 (Tyr705) (CST 9145, RRID: AB_2491009), in 5% BSA-TBST was incubated with membranes overnight at 4°C. 1:1000 STAT3 (CST 9139, RRID: AB_3528142) primary antibody was incubated with membranes overnight in 5% Milk TBS-T. Following TBS-T washes, 1:3000 secondary antibody anti-rabbit (Cell Signaling Technology IgG, HRP linked #7074, RRID: AB_2099233) in 5% BSA-TBST for pSTAT3 and 5% Milk TBS-T for STAT3 was added for 1 hour rocking at RT. After TBS-T washes, membranes were quickly washed with 1 x TBS wash and developed using SuperSignal™ West Femto Maximum Sensitivity Substrate (Thermo Fisher Scientific #34094) and Clarity Western ECL substrate (Bio-Rad #1705061) as per manufacturer’s protocol with the Azure 600 Imager (Azure Biosystems) for imaging at autoexposure. Band volumes for pSTAT3 and STAT3 were quantified using AzureSpot Pro analysis or ImageJ. Band volume for pSTAT3 was divided by STAT3 for fold change. Two to four independent experiments were conducted. SimplyBlue SafeStain (Invitrogen^TM^ #LC6060) was utilized to stain membrane for evaluation of equal loading and storage in binder for scanning, storage, and documentation of the membranes.

#### For cells that were stimulated

MCF-7 and T-47D cells were cultured and seeded in 6 well plates using their respective complete media. Once 60-80% cell confluence was reached in 6 well plates, cells were serum starved overnight for a minimum of 18 hours and a maximum of 24 hours. Cells were then incubated with the following either alone or in combination in DMEM SFM (control media): 1) DMEM SFM – 1 to 2 mL of DMEM SFM, 2) HBVP CM – 1 to 2 mL of pericyte conditioned media from a T-75 flask, 3) vehicle-treated HBVP CM, 4) 10 nM DTX-treated HBVP CM, 5) 0.1 µg/mL control IgG, 6) 0.1 µg/mL tocilizumab, 7) 10 nM DTX, 8) vehicle, 9) 2.5 ng/mL IL-6, and 10) 5.0 ng/mL IL-6. Following stimulation, cells were washed 1x with ice-cold PBS. The same protocol as above was followed post-cell harvest.

### Statistical analysis

All data were analyzed in either GraphPad Prism 10 (Boston, MA) or using R Studio. All flow cytometry data were analyzed using FlowJo v10 or floreada.io. Results were analyzed as Mean ± s.e.m. as noted, accordingly. The *P* values in these experiments were determined based on biological and not technical replicates. Statistical tests are explicitly included in the results and figure legends.

#### Organoid growth modeling

Growth rates were estimated by fitting a linear model to log-transformed area measurements as a function of time, condition, and their interaction. The data were fit globally to constrain a single estimate of A_0_. Condition-specific growth rates were extracted as the slopes of time using estimated marginal trends. Differences in growth rates were assessed using an ANCOVA model with Bonferroni correction for multiple comparisons.

#### Bliss excess

Drug interactions were quantified using a Bliss independence framework adapted for exponential growth dynamics. Growth rates (*k*) were estimated by fitting an exponential model to longitudinal culture area measurements for each condition. Relative effects at time *t* were computed as

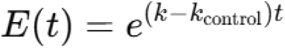

where *k*_control_ corresponds to the matched vehicle control. Bliss excess was defined as

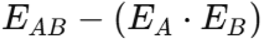

with negative values indicating synergistic growth inhibition and positive values indicating antagonism. Confidence bounds were derived by propagating uncertainty from the fitted growth-rate estimates; uncertainty in the control condition was not propagated.

All organoid analyses were performed in RStudio (version 2024.09.1+394) and R (version 4.4.1).

## Data and code availability

Research outputs will be preserved and shared with the public via Knowledge@UChicago, the University of Chicago institutional repository. Knowledge@UChicago is managed by the University Library and uses a cloud-based hosted repository service called TIND as its platform. TIND is based on the Invenio software originally developed at the research institute CERN to manage its own scholarly output.

TIND uses an OAIS-compliant approach to preservation, with redundancy backup, conversion to archival file formats, storage of archival packages on geographically separated servers, and fixity checking. To support discoverability and citation, DOIs are minted for deposits made to Knowledge@UChicago through the DataCite Fabrica service.

All data generated or analyzed during this study are included in the Article or as Supplemental Information. The source data underlying selected Figures and Supplementary Figures are provided as a Source Data file. The data underlying the findings of this study can be obtained from the corresponding author upon reasonable request. Please contact Samantha S. Yee at.

## Acknowledgements

We would like to thank Jinyang Li, Anthony Kossiakoff, and Marsha Rosner for cell lines. Thank you to the University of Chicago Cytometry and Antibody Technology (CAT) Facility (RRID: SCR_017760). We appreciate the Janes’ Laboratory for the feedback provided on the figures. Also, we thank Charles Fermaintt, Sridevi Challa, Kevin Muñoz Forti, and Carrie W. Rinker-Schaeffer for their scientific discussions and insight. Also, we thank Jessica J. Kandel for her continuous support. We sincerely thank Kevin A. Janes and Shayna L. Showalter for helping with clinical material procurement for this study, Alexys Riddick for the EdU protocol and microscope assistance, and Janes’ laboratory members for critical feedback.

## Author contributions

Conceptualization: S.S.Y.; Methodology: S.S.Y., and R.K.P.; Investigation: S.S.Y., R.K.P., J.G., and V.J.L.; Formal analysis: S.S.Y., R.K.P., J.G., and V.J.L.; Writing—original draft: S.S.Y., and R.K.P; Writing—review and editing: S.S.Y., R.K.P., S.L.H., V.J.L., J.G., N.A.J., and L.V. All authors reviewed and approved the manuscript. Funding Acquisition: S.S.Y. and R.K.P.; Supervision: S.S.Y.; Project administration: S.S.Y.

## Ethics declarations (approval and consent to participate)

Human sample acquisition and experimental procedures were carried out in compliance with regulations and protocols approved by the Institutional Review Board for Health Sciences Research (IRB-HSR) at the University of Virginia in accordance with the U.S. Common Rule and IRB Protocol #14176. The Institutional Review Board has granted this study a waiver of consent under 45CFR46.116 of the 2018 Common Rule.

## Additional information

We have used the following abbreviations

## Introduction/Results/Discussion

DTX: docetaxel
HBVP: human brain vascular pericytes, pericytes
MTA: microtubule targeting agent
ER+: estrogen receptor positive
IL-6/JAK/STAT3: Interleukin-6/Janus kinase/Signal transducer and activator of transcription 3
pSTAT3: phosphorylated STAT3
TME: tumor microenvironment
IL-6R: interleukin-6 receptor
IL-6Rα: alpha subunit
IL-6Rβ: beta subunit
HER2: human epidermal growth factor receptor 2
JAK2: Janus kinase 2
IGF-2: insulin-like growth factor 2
TGF-β: transforming growth factor beta
CXCL1: C-X-C Motif Chemokine Ligand 1
CXCL5: C-X-C Motif Chemokine Ligand 5
ST2: suppression of tumorigenicity 2
GDF-15: growth differentiation factor 15
ELISA: enzyme-linked immunosorbent assay
RT-qPCR: real-time quantitative polymerase chain reaction
GAPDH: glyceraldehyde 3-phosphate dehydrogenase
TNBC: triple-negative breast cancer
CM: conditioned media
HBVP CM: human brain vascular pericyte conditioned media
sIL-6Rα: soluble IL6-Rα
mIL-6Rα: membrane-bound IL-6Rα
IgG: immunoglobulin G
TLR4: Toll-like receptor 4
G-CSF: granulocyte-colony stimulating factor
TNFα: tumor necrosis factor α
NF-κB: nuclear factor kappa-light-chain-enhancer of activated B cells
MD-2: myeloid differentiation 2
LPS: lipopolysaccharide
DAPI: 4′,6-diamidino-2-phenylindole
EdU: 5-ethynyl-2’-deoxyuridine

## Methods

DMSO: dimethyl sulfoxide
HRP-linked: horseradish peroxidase-linked
CD126: cluster of differentiation 126
CD130: glycoprotein 130
ATCC: American Type Culture Collection
FBS: fetal bovine serum
DMEM: Dulbecco’s Modified Eagle Medium
P/S: penicillin-streptomycin
RPMI: Roswell Park Memorial Institute
IMEM: Improved Minimum Essential Medium
BSA: bovine serum albumin
ROCK inhibitor: Rho kinase inhibitor
RBC lysis buffer: red blood cell lysis buffer
PBS: phosphate-buffered saline
SFM: serum-free media
trypsin-EDTA: trypsin-ethylenediaminetetraacetic acid
APC (flow cytometry): allophycocyanin
PE (flow cytometry): phycoerythrin
UNG incubation: Uracil-N-Glycosylase incubation
cDNA: complementary DNA
CCK8: cell counting kit-8
RIPA buffer: RadioImmunoprecipitation Assay buffer
LDS buffer: lithium dodecyl sulfate buffer
PVDF: polyvinylidene difluoride
TBS: Tris-Buffered Saline
TBS-T: Tris-Buffered Saline with Tween-20
RT: room temperature
ECL substrate: Enhanced Chemiluminescence substrate
NA: numerical aperture
ANCOVA model: analysis of covariance model

## Supplementary Figures and Tables

**Supplementary Figure 1.**
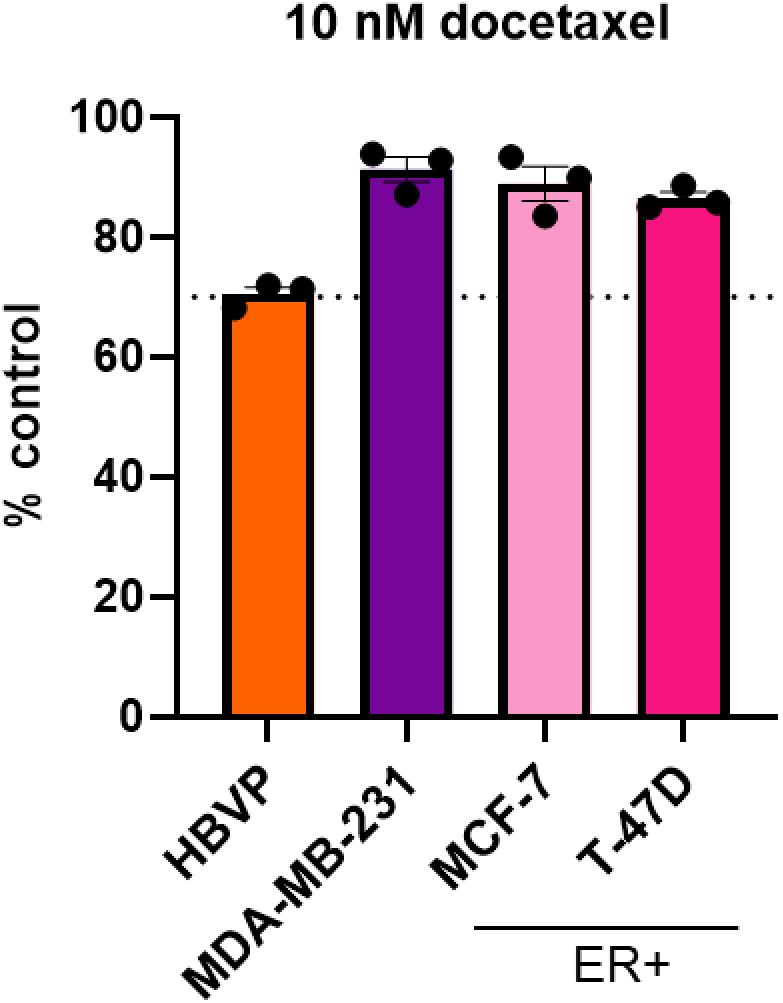
A minimum of 70% cell viability (% control) is achieved in pericytes and 3 breast cancer cell lines following 24-hour treatment of 10 nM docetaxel via CCK-8 assay. Dashed line indicated at 70%. Data is representative of n = 3 independent experiments. Mean ± s.e.m. shown. HBVP = human brain vascular pericytes.

**Supplementary Figure 2.**
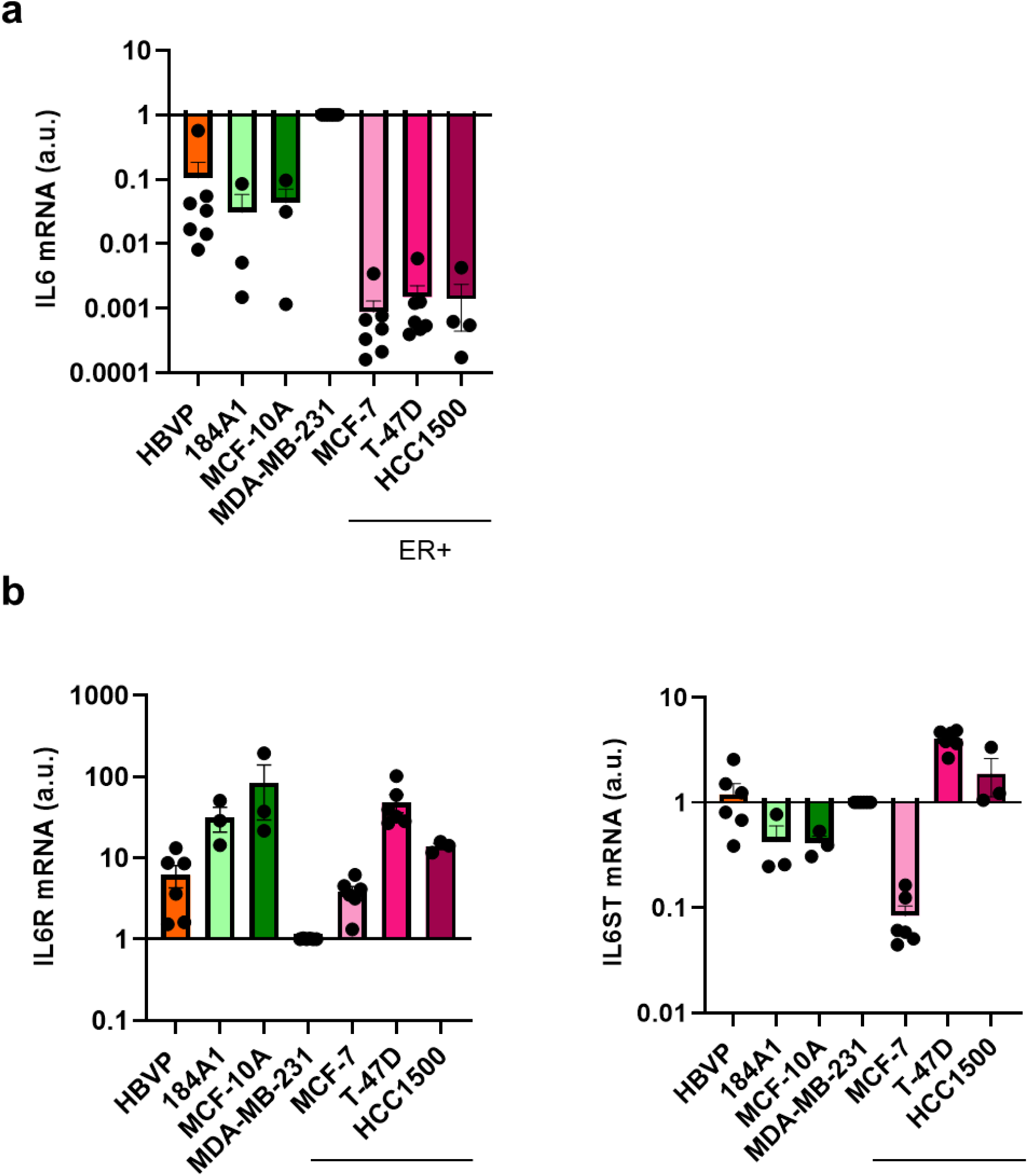
Pericytes, non-tumorigenic breast epithelial cell lines, and breast cancer cell lines have varying basal levels of (a) *IL6*, (b) *IL6R* and *IL6ST* mRNA expression. **a** *IL6* mRNA expression of untreated cells and cell lines in reference to *GAPDH*. a.u. denotes arbitrary units. Data is representative of n = 4 - 7 independent experiments. Mean ± s.e.m. shown. **b** Basal *IL6R* (left) and *IL6ST* (right) transcript levels using reference gene *GAPDH* relative to the MDA-MB-231 cell line normalized to 1. a.u. = arbitrary units. Data is representative of n = 3 - 6 independent experiments with technical duplicates. Mean ± s.e.m. shown. HBVP = human brain vascular pericytes.

**Supplementary Figure 3.**
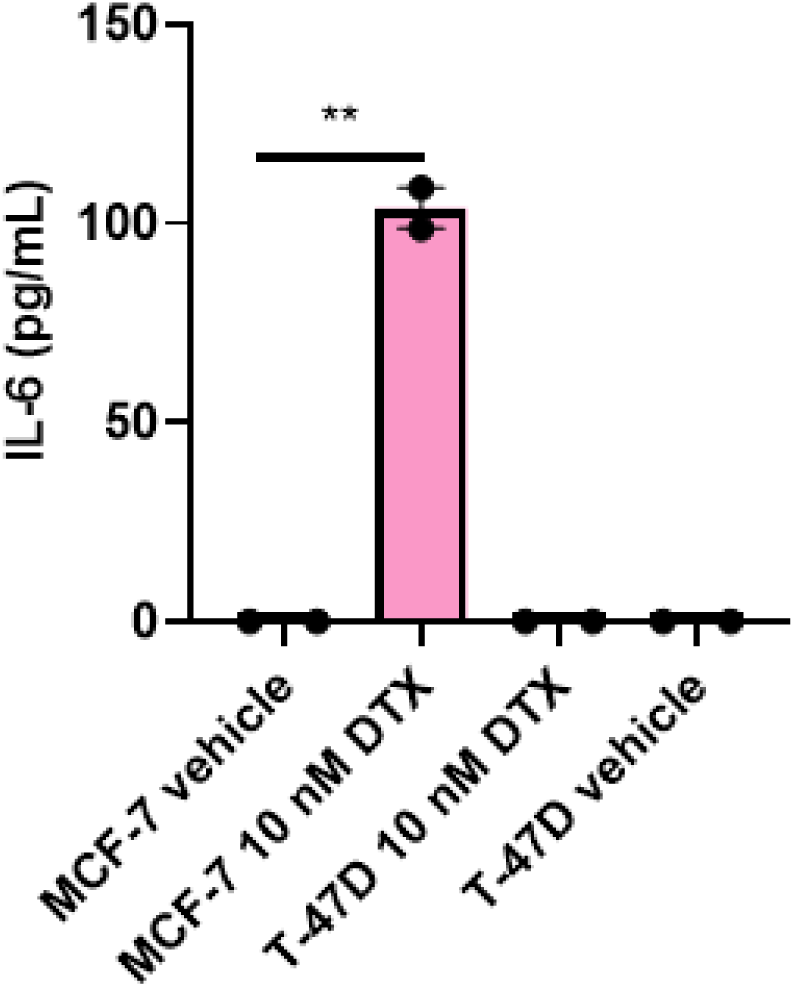
Secreted IL-6 from docetaxel-treated ER+ breast cell lines. IL-6 (pg/mL) quantified using conditioned media from vehicle- or 10 nM DTX-treated MCF-7 and T-47D cell lines following 24-hour treatment via an IL-6 ELISA. Data is representative of n = 2 independent experiments with technical duplicates. Mean ± s.e.m. shown. *P* values were calculated using two-tailed unpaired t tests. ** *P* ≤ 0.01. DTX = docetaxel.

**Supplementary Figure 4.**
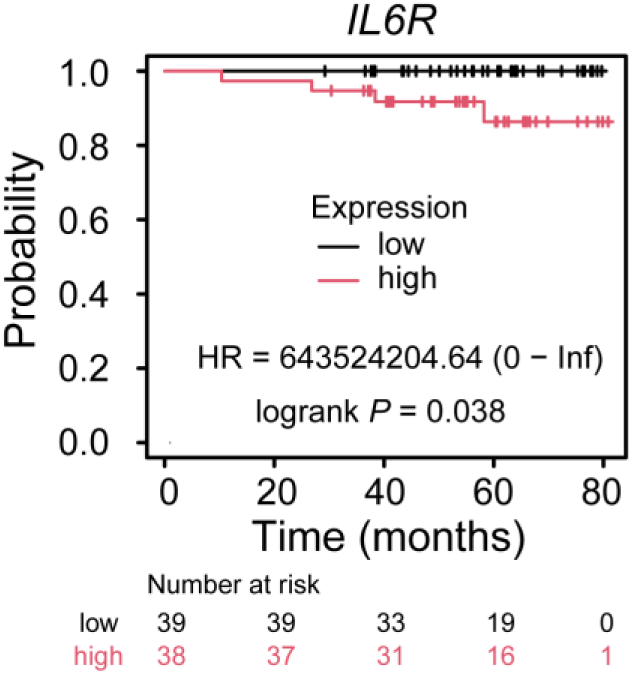
**High *IL6R* expression (Q4) is associated with worse overall survival in luminal A (ER+/HER2-) treatment-naïve breast cancer patients (n=155).**

**Supplementary Figure 5.**
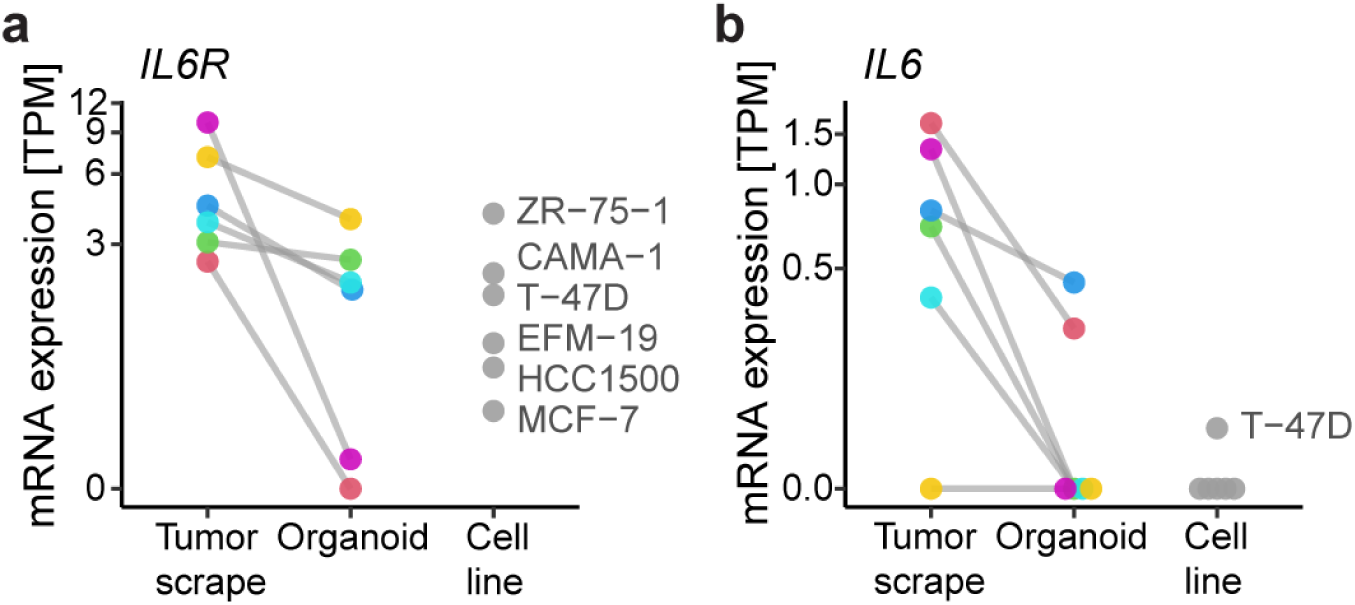
**Zero-passage patient-derived ER+ breast cancer organoids preserve *IL6R* expression in most cases (a) and *IL6* expression in a subset (b) in a non-exclusive way.**

**Supplementary Table 1.** Characterization of cells used in this study.

| **Human cell name** | **Cancer type** | **Primary cells or cell line** | **P53 status** | **Origin of cells /Molecular subtypes** | **Race/ethnicity and other information** | **References** |
| --- | --- | --- | --- | --- | --- | --- |
| Human brain vascular pericytes (HBVP) | Non-cancerous | Primary cells | Not available | Not applicable | Female donor, gestational age 22 weeks | ScienCell (#1201) |
| MDA-MB-231 | TNBC | Cell line | Mutant | Metastatic, pleural effusion | White | ATCC (# HTB-26), DepMap ID: ACH-000768<sup>122</sup> |
| HCC1500 | ER+ breast | Cell line | Wild-type | Primary - breast | Black | ATCC (#CRL-2329), and DepMap ID: ACH-000349 |
| MCF-7 | ER+ breast | Cell line | Wild-type | Metastatic, pleural effusion | White | ATCC (#HTB-22), and DepMap ID: ACH-000019 |
| T-47D | ER+ breast | Cell line | Mutant | Metastatic, pleural effusion | White | ATCC (#HTB-133), DepMap ID: ACH-000147<sup>122</sup> |
| MCF-10A | Non-cancerous: Breast - epithelial | Non-tumorigenic epithelial cell line | Wild-type | Primary - breast | White | ATCC (#CRL-10317) and DepMap ID: ACH-001357 |
| 184A1 | Non-cancerous: Breast epithelial-like | Non-tumorigenic epithelial-like cell | Wild-type | Derived from mammary gland tissue | Not available | ATCC (#CRL-8798)<sup>123</sup> |

**Supplementary Table 2.** Patient characteristics for ER+ breast tumor organoids.

| Case | UVABCO176 | UVABCO178 | UVABCO179 |
| --- | --- | --- | --- |
| Sex | Female | Female | Female |
| Age (years) | 51 | 59 | 34 |
| Race | White | White | White |
| Ethnicity | Non-Hispanic | Non-Hispanic | Non-Hispanic |
| Laterality | Right | Left | Left |
| Surgery_Type | Lumpectomy | Mastectomy | Mastectomy |
| Dx_Biopsy | Invasive ductal carcinoma | Invasive ductal carcinoma with lobular features | Invasive ductal carcinoma |
| Grade_Biopsy | 1 | 2 | 2 |
| ER_Biopsy | 3+ | 3+ | 3+ |
| ER_Percentage | 95% | 95% | 80% |
| PR_Biopsy | 3+ | 3+ | NA |
| PR_Percentage | 95% | 75% | NA |
| HER2_Biopsy | Negative | Negative | Negative |
| Dx_Resection | Invasive ductal carcinoma with lobular features | Invasive ductal carcinoma | Invasive ductal carcinoma |
| Grade_Resection | 1 | 2 | 3 |
| Size_Resection_mm | 9 | 55 | 25 |
| Stage_Resection | pT1bN0sn | pT3N1asn | pT2mN1a |
| Size_Imaging_mm | 7 | 45 | 25 |

Abbreviations: NA = Not Available

## References

1. Siegel, R. L., Kratzer, T. B., Wagle, N. S., Sung, H. & Jemal, A. Cancer statistics, 2026. CA. Cancer J. Clin. 76, e70043 (2026).

2. Surveillance Research Program, National Cancer Institute. SEER*Explorer: An interactive website for SEER cancer statistics [Internet]. SEER Incidence Data, November 2024 Submission (1975-2022), SEER 21 registries. (2025).

3. Ring, A. & Dowsett, M. Mechanisms of tamoxifen resistance. Endocr. Relat. Cancer 11, 643–658 (2004).

4. van Maaren, M. C. et al. Ten-year recurrence rates for breast cancer subtypes in the Netherlands: A large population-based study. Int. J. Cancer 144, 263–272 (2019).

5. Gallicchio, L., Devasia, T. P., Tonorezos, E., Mollica, M. A. & Mariotto, A. Estimation of the Number of Individuals Living With Metastatic Cancer in the United States. JNCI J. Natl. Cancer Inst. 114, 1476–1483 (2022).

6. Trudeau, M. E. Docetaxel (Taxotere): an overview of first-line monotherapy. Semin. Oncol. 22, 17–21 (1995).

7. Postigo-Corrales, F. et al. Docetaxel Resistance in Breast Cancer: Current Insights and Future Directions. Int. J. Mol. Sci. 26, (2025).

8. Andre, F. et al. Estrogen Receptor Expression and Docetaxel Efficacy in Patients with Metastatic Breast Cancer: A Pooled Analysis of Four Randomized Trials. The Oncologist 15, 476–483 (2010).

9. O’Shaughnessy, J. Extending Survival with Chemotherapy in Metastatic Breast Cancer. The Oncologist 10, 20–29 (2005).

10. Rivera, E. & Gomez, H. Chemotherapy resistance in metastatic breast cancer: the evolving role of ixabepilone. Breast Cancer Res. 12, S2 (2010).

11. Dhiman, V. K., Kumari, M. & Singh, D. Chemoresistance: The hidden barrier in cancer treatment. Cancer Pathog. Ther. 4, 98–109 (2026).

12. Liang, Y., Shao, Y. & Gu, W. Role of interleukin-6 in resistance to tumor therapy. Discov. Oncol. 16, 1791 (2025).

13. Wang, H. et al. Crosstalk of pyroptosis and cytokine in the tumor microenvironment: from mechanisms to clinical implication. Mol. Cancer 23, 268 (2024).

14. Lee, H.-J. et al. Drug Resistance via Feedback Activation of Stat3 in Oncogene-Addicted Cancer Cells. Cancer Cell 26, 207–221 (2014).

15. Sreenivasan, L. et al. Autocrine IL-6/STAT3 signaling aids development of acquired drug resistance in Group 3 medulloblastoma. Cell Death Dis. 11, 1035 (2020).

16. Sen, M. et al. Targeting Stat3 Abrogates EGFR Inhibitor Resistance in Cancer. Clin. Cancer Res. 18, 4986–4996 (2012).

17. Sabaawy, H. E., Ryan, B. M., Khiabanian, H. & Pine, S. R. JAK/STAT of all trades: linking inflammation with cancer development, tumor progression and therapy resistance. Carcinogenesis 42, 1411–1419 (2021).

18. Bromberg, J. F. et al. Stat3 as an Oncogene. Cell 98, 295–303 (1999).

19. Zhang, T. & Xiaohan, C. Unveiling the Role of JAK2/STAT3 signaling in chemoresistance of gynecological cancers: From mechanisms to therapeutic implications. Crit. Rev. Oncol. Hematol. 211, 104712 (2025).

20. Garcia, R. et al. Constitutive activation of Stat3 by the Src and JAK tyrosine kinases participates in growth regulation of human breast carcinoma cells. Oncogene 20, 2499–2513 (2001).

21. Wischnewski, V. et al. Phenotypic diversity of T cells in human primary and metastatic brain tumors revealed by multiomic interrogation. *Nat*. Cancer 4, 908–924 (2023).

22. Visser, K. E. de & Joyce, J. A. The evolving tumor microenvironment: From cancer initiation to metastatic outgrowth. Cancer Cell 41, 374–403 (2023).

23. Anderson, N. M. & Simon, M. C. The tumor microenvironment. Curr. Biol. 30, R921–R925 (2020).

24. Ganesan, R. et al. Taxane chemotherapy induces stromal injury that leads to breast cancer dormancy escape. PLOS Biol. 21, e3002275 (2023).

25. Miroshnychenko, D. et al. Stroma-Mediated Breast Cancer Cell Proliferation Indirectly Drives Chemoresistance by Accelerating Tumor Recovery between Chemotherapy Cycles. Cancer Res. 83, 3681–3692 (2023).

26. Skandha Gopalan, K. & Bergers, G. The Pan-Tumor Vasculature under the Transcriptomic Magnifying Glass. Cancer Res. 84, 3502–3504 (2024).

27. Pan, X. et al. Tumour vasculature at single-cell resolution. Nature 632, 429–436 (2024).

28. Lyle, L. T. et al. Alterations in Pericyte Subpopulations Are Associated with Elevated Blood–Tumor Barrier Permeability in Experimental Brain Metastasis of Breast Cancer. Clin. Cancer Res. 22, 5287–5299 (2016).

29. Fouladzadeh, A. et al. The development of tumour vascular networks. *Commun*. Biol. 4, 1111 (2021).

30. Stratman, A. N. & Davis, G. E. Endothelial Cell-Pericyte Interactions Stimulate Basement Membrane Matrix Assembly: Influence on Vascular Tube Remodeling, Maturation, and Stabilization. Microsc. Microanal. 18, 68–80 (2012).

31. Morikawa, S. et al. Abnormalities in Pericytes on Blood Vessels and Endothelial Sprouts in Tumors. Am. J. Pathol. 160, 985–1000 (2002).

32. Cuevas, P. et al. Pericyte endothelial gap junctions in human cerebral capillaries. Anat. Embryol. (Berl.) 170, 155–159 (1984).

33. Birbrair, A. Pericyte Biology: Development, Homeostasis, and Disease. in Pericyte Biology - Novel Concepts (ed. Birbrair, A.) 1–3 (Springer International Publishing, Cham, 2018). doi:10.1007/978-3-030-02601-1_1.

34. Jordan, A. M., Drake, R. J. G. & Hodivala-Dilke, K. M. Angiocrine and pericrine signaling: how endothelial cells and pericytes drive cancer progression and therapy resistance. Physiol. Rev. 106, 87–119 (2026).

35. Naumov, G. N., Folkman, J., Straume, O. & Akslen, L. A. Tumor-vascular interactions and tumor dormancy. APMIS 116, 569–585 (2008).

36. Carmeliet, P. Angiogenesis in health and disease. Nat. Med. 9, 653–660 (2003).

37. Bose, A. et al. Tumor-Derived Vascular Pericytes Anergize Th Cells. J. Immunol. 191, 971–981 (2013).

38. Chen, M. et al. Pericyte-targeting prodrug overcomes tumor resistance to vascular disrupting agents. J. Clin. Invest. 127, 3689–3701 (2017).

39. Abramsson, A., Lindblom, P. & Betsholtz, C. Endothelial and nonendothelial sources of PDGF-B regulate pericyte recruitment and influence vascular pattern formation in tumors. J. Clin. Invest. 112, 1142–1151 (2003).

40. Del Toro, K. et al. Breast pericytes: a newly identified driver of tumor cell proliferation. Front. Oncol. 14, (2024).

41. Barron, L., Gharib, S. A. & Duffield, J. S. Lung Pericytes and Resident Fibroblasts: Busy Multitaskers. Am. J. Pathol. 186, 2519–2531 (2016).

42. Gaceb, A., Barbariga, M., Özen, I. & Paul, G. The pericyte secretome: Potential impact on regeneration. Biochimie 155, 16–25 (2018).

43. Molnár, K. et al. Pericyte-secreted IGF2 promotes breast cancer brain metastasis formation. Mol. Oncol. 14, 2040–2057 (2020).

44. Walshe, T. E. et al. TGF-β Is Required for Vascular Barrier Function, Endothelial Survival and Homeostasis of the Adult Microvasculature. PLOS ONE 4, e5149 (2009).

45. Shimizu, F. et al. Advanced glycation end-products disrupt the blood–brain barrier by stimulating the release of transforming growth factor–β by pericytes and vascular endothelial growth factor and matrix metalloproteinase–2 by endothelial cells in vitro. Neurobiol. Aging 34, 1902–1912 (2013).

46. Moro, M., Balestrero, F. C. & Grolla, A. A. Pericytes: jack-of-all-trades in cancer-related inflammation. Front. Pharmacol. 15, (2024).

47. Li, X. et al. TCAF2 in Pericytes Promotes Colorectal Cancer Liver Metastasis via Inhibiting Cold-Sensing TRPM8 Channel. Adv. Sci. 10, 2302717 (2023).

48. Prete, A. et al. Pericytes Elicit Resistance to Vemurafenib and Sorafenib Therapy in Thyroid Carcinoma via the TSP-1/TGFβ1 Axis. Clin. Cancer Res. 24, 6078–6097 (2018).

49. McErlain, T. et al. Pericyte-tumor crosstalk facilitates metastatic tumor cell latency through PIEZO1-activated lysophospholipid transfer. BioRxiv Prepr. Serv. Biol. 2025.02.15.638347 (2025) doi:10.1101/2025.02.15.638347.

50. Ghajar, C. M. et al. The perivascular niche regulates breast tumour dormancy. Nat. Cell Biol. 15, 807–817 (2013).

51. Gaceb, A. & Paul, G. Pericyte Secretome. in Pericyte Biology - Novel Concepts (ed. Birbrair, A.) 139–163 (Springer International Publishing, Cham, 2018). doi:10.1007/978-3-030-02601-1_11.

52. Kovac, A., Erickson, M. A. & Banks, W. A. Brain microvascular pericytes are immunoactive in culture: cytokine, chemokine, nitric oxide, and LRP-1 expression in response to lipopolysaccharide. J. Neuroinflammation 8, 139 (2011).

53. Nehmé, A. & Edelman, J. Dexamethasone Inhibits High Glucose–, TNF-α–, and IL-1β–Induced Secretion of Inflammatory and Angiogenic Mediators from Retinal Microvascular Pericytes. Invest. Ophthalmol. Vis. Sci. 49, 2030–2038 (2008).

54. Pieper, C., Pieloch, P. & Galla, H.-J. Pericytes support neutrophil transmigration via interleukin-8 across a porcine co-culture model of the blood–brain barrier. Brain Res. 1524, 1–11 (2013).

55. Baker, S. D. et al. Comparative Pharmacokinetics of Weekly and Every-Three-Weeks Docetaxel. Clin. Cancer Res. 10, 1976–1983 (2004).

56. Verweij, J., Clavel, M. & Chevalier, B. Paclitaxel (TaxolTM) and docetaxel (TaxotereTM): Not simply two of a kind. Ann. Oncol. 5, 495–505 (1994).

57. Grant, D. S., Williams, T. L., Zahaczewsky, M. & Dicker, A. P. Comparison of antiangiogenic activities using paclitaxel (taxol) and docetaxel (taxotere). Int. J. Cancer 104, 121–129 (2003).

58. Vacca, A. et al. Docetaxel versus paclitaxel for antiangiogenesis. J. Hematother. Stem Cell Res. 11, 103–118 (2002).

59. Iida, S., Shimada, J. & Sakagami, H. Cytotoxicity Induced by Docetaxel in Human Oral Squamous Cell Carcinoma Cell Lines. In Vivo 27, 321–332 (2013).

60. Fermaintt, C. S., Takahashi-Ruiz, L., Liang, H., Mooberry, S. L. & Risinger, A. L. Eribulin Activates the cGAS-STING Pathway via the Cytoplasmic Accumulation of Mitochondrial DNA. Mol. Pharmacol. 100, 309–318 (2021).

61. Wang, N. et al. CXCL1 derived from tumor-associated macrophages promotes breast cancer metastasis via activating NF-κB/SOX4 signaling. Cell Death Dis. 9, 880 (2018).

62. Mao, Z. et al. CXCL5 promotes gastric cancer metastasis by inducing epithelial-mesenchymal transition and activating neutrophils. Oncogenesis 9, 63 (2020).

63. Lu, D. et al. Serum soluble ST2 is associated with ER-positive breast cancer. BMC Cancer 14, 198 (2014).

64. Fang, M. et al. Functional characteristics of fresh antitumor immune interferer GDF-15 in multiple cancers. Sci. Rep. 15, 30864 (2025).

65. Lee, S. O. et al. Interleukin-6 promotes androgen-independent growth in LNCaP human prostate cancer cells. Clin. Cancer Res. Off. J. Am. Assoc. Cancer Res. 9, 370–376 (2003).

66. Chang, Q. et al. The IL-6/JAK/Stat3 Feed-Forward Loop Drives Tumorigenesis and Metastasis. Neoplasia 15, 848-IN45 (2013).

67. Said, E. A. et al. Defining IL-6 levels in healthy individuals: A meta-analysis. J. Med. Virol. 93, 3915–3924 (2021).

68. Lokau, J. et al. Long-term increase in soluble interleukin-6 receptor levels in convalescents after mild COVID-19 infection. Front. Immunol. 15, (2025).

69. Baran, P. et al. The balance of interleukin (IL)-6, IL-6·soluble IL-6 receptor (sIL-6R), and IL-6·sIL-6R·sgp130 complexes allows simultaneous classic and trans-signaling. J. Biol. Chem. 293, 6762–6775 (2018).

70. Tanaka, T., Narazaki, M. & Kishimoto, T. IL-6 in Inflammation, Immunity, and Disease. Cold Spring Harb. Perspect. Biol. 6, a016295 (2014).

71. Sasser, A. K. et al. Interleukin-6 is a potent growth factor for ER-α-positive human breast cancer. FASEB J. 21, 3763–3770 (2007).

72. Chen, J. et al. IL-6: The Link Between Inflammation, Immunity and Breast Cancer. Front. Oncol. 12, (2022).

73. Siersbæk, R. et al. IL6/STAT3 Signaling Hijacks Estrogen Receptor α Enhancers to Drive Breast Cancer Metastasis. Cancer Cell 38, 412–423.e9 (2020).

74. Yamasaki, K. et al. Cloning and Expression of the Human Interleukin-6 (BSF-2/IFNβ 2) Receptor. Science 241, 825–828 (1988).

75. Mosly, D. et al. Variation in IL6ST cytokine family function and the potential of IL6 trans-signalling in ERα positive breast cancer cells. Cell. Signal. 103, 110563 (2023).

76. Tsoi, H., Man, E. P. S., Chau, K. M. & Khoo, U.-S. Targeting the IL-6/STAT3 Signalling Cascade to Reverse Tamoxifen Resistance in Estrogen Receptor Positive Breast Cancer. Cancers 13, (2021).

77. Kang, S. & Kishimoto, T. Interplay between interleukin-6 signaling and the vascular endothelium in cytokine storms. Exp. Mol. Med. 53, 1116–1123 (2021).

78. Mihara, M. et al. Tocilizumab inhibits signal transduction mediated by both mIL-6R and sIL-6R, but not by the receptors of other members of IL-6 cytokine family. Int. Immunopharmacol. 5, 1731–1740 (2005).

79. Chen, L. Y. C. et al. Soluble interleukin-6 receptor in the COVID-19 cytokine storm syndrome. Cell Rep. Med. 2, 100269 (2021).

80. Xue, C. et al. Evolving cognition of the JAK-STAT signaling pathway: autoimmune disorders and cancer. Signal Transduct. Target. Ther. 8, 204 (2023).

81. Győrffy, B. Survival analysis across the entire transcriptome identifies biomarkers with the highest prognostic power in breast cancer. Comput. Struct. Biotechnol. J. 19, 4101–4109 (2021).

82. Przanowska, R. K. et al. Patient-derived response estimates from zero-passage organoids of luminal breast cancer. Breast Cancer Res. 26, 192 (2024).

83. Flobak, Å. et al. A high-throughput drug combination screen of targeted small molecule inhibitors in cancer cell lines. Sci. Data 6, 237 (2019).

84. Reck, M. et al. Docetaxel plus nintedanib versus docetaxel plus placebo in patients with previously treated non-small-cell lung cancer (LUME-Lung 1): a phase 3, double-blind, randomised controlled trial. Lancet Oncol. 15, 143–155 (2014).

85. Richeldi, L. et al. Efficacy and Safety of Nintedanib in Idiopathic Pulmonary Fibrosis. N. Engl. J. Med. 370, 2071–2082 (2014).

86. Early Breast Cancer Trialists’ Collaborative Group (EBCTCG). Effects of chemotherapy and hormonal therapy for early breast cancer on recurrence and 15-year survival: an overview of the randomised trials. Lancet 365, 1687–1717 (2005).

87. Demicheli, R., Abbattista, A., Miceli, R., Valagussa, P. & Bonadonna, G. Time distribution of the recurrence risk for breast cancer patients undergoing mastectomy: further support about the concept of tumor dormancy. Breast Cancer Res. Treat. 41, 177–185 (1996).

88. Karrison, T. G., Ferguson, D. J. & Meier, P. Dormancy of Mammary Carcinoma After Mastectomy. JNCI J. Natl. Cancer Inst. 91, 80–85 (1999).

89. Fernandes Neto, J. M., et al. Multiple low dose therapy as an effective strategy to treat EGFR inhibitor-resistant NSCLC tumours. Nat. Commun. 11, 3157 (2020).

90. Chan, T.-S. et al. Metronomic chemotherapy prevents therapy-induced stromal activation and induction of tumor-initiating cells. J. Exp. Med. 213, 2967–2988 (2016).

91. França, G. S. et al. Cellular adaptation to cancer therapy along a resistance continuum. Nature 631, 876–883 (2024).

92. Wang, Z. et al. Drug-tolerant persister cells in cancer: bridging the gaps between bench and bedside. Nat. Commun. 16, 10048 (2025).

93. Rose-John, S. Interleukin-6 Family Cytokines. Cold Spring Harb. Perspect. Biol. 10, a028415 (2018).

94. Lai, C.-F. et al. Receptors for Interleukin (IL)-10 and IL-6-type Cytokines Use Similar Signaling Mechanisms for Inducing Transcription through IL-6 Response Elements *. J. Biol. Chem. 271, 13968–13975 (1996).

95. Leung, E. et al. Pharmacokinetic/Pharmacodynamic Considerations of Alternate Dosing Strategies of Tocilizumab in COVID-19. Clin. Pharmacokinet. 61, 155–165 (2022).

96. Burmester, G. R. et al. A randomised, double-blind, parallel-group study of the safety and efficacy of subcutaneous tocilizumab versus intravenous tocilizumab in combination with traditional disease-modifying antirheumatic drugs in patients with moderate to severe rheumatoid arthritis (SUMMACTA study). Ann. Rheum. Dis. 73, 69–74 (2014).

97. Hibi, M. et al. Molecular cloning and expression of an IL-6 signal transducer, gp130. Cell 63, 1149–1157 (1990).

98. Müllberg, J. et al. Differential shedding of the two subunits of the interleukin-6 receptor. FEBS Lett. 332, 174–178 (1993).

99. Riethmueller, S. et al. Proteolytic Origin of the Soluble Human IL-6R In Vivo and a Decisive Role of N-Glycosylation. PLOS Biol. 15, e2000080 (2017).

100. Osuala, K. O. et al. Il-6 signaling between ductal carcinoma in situ cells and carcinoma-associated fibroblasts mediates tumor cell growth and migration. BMC Cancer 15, 584 (2015).

101. Chung, A. W. et al. Tocilizumab overcomes chemotherapy resistance in mesenchymal stem-like breast cancer by negating autocrine IL-1A induction of IL-6. Npj Breast Cancer 8, 30 (2022).

102. Zhang, N., Fu, J.-N. & Chou, T.-C. Synergistic combination of microtubule targeting anticancer fludelone with cytoprotective panaxytriol derived from panax ginseng against MX-1 cells in vitro: experimental design and data analysis using the combination index method. Am. J. Cancer Res. 6, 97–104 (2016).

103. Chou, T.-C. Drug Combination Studies and Their Synergy Quantification Using the Chou-Talalay Method. Cancer Res. 70, 440–446 (2010).

104. Alraouji, N. N. & Aboussekhra, A. Tocilizumab inhibits IL-8 and the proangiogenic potential of triple negative breast cancer cells. Mol. Carcinog. 60, 51–59 (2021).

105. Pederson, P. J., Liang, H., Filonov, D. & Mooberry, S. L. Eribulin and Paclitaxel Differentially Alter Extracellular Vesicles and Their Cargo from Triple-Negative Breast Cancer Cells. Cancers 13, (2021).

106. Hassouneh, Z. et al. Low-Dose Eribulin Promotes NK Cell-Mediated Therapeutic Efficacy in Bladder Cancer. Cancers 16, (2024).

107. Kaul, R., Risinger, A. L. & Mooberry, S. L. Microtubule-Targeting Drugs: More than Antimitotics. J. Nat. Prod. 82, 680–685 (2019).

108. Ganesan, R. et al. Taxane chemotherapy induces stromal injury that leads to breast cancer dormancy escape. PLOS Biol. 21, e3002275 (2023).

109. Edwardson, D. W. et al. Inflammatory cytokine production in tumor cells upon chemotherapy drug exposure or upon selection for drug resistance. PLOS ONE 12, e0183662 (2017).

110. Liu, L., Liu, Y., Yan, X., Zhou, C. & Xiong, X. The role of granulocyte colony-stimulating factor in breast cancer development: A review. Mol. Med. Rep. 21, 2019–2029 (2020).

111. Sprowl, J. A. et al. Alterations in tumor necrosis factor signaling pathways are associated with cytotoxicity and resistance to taxanes: a study in isogenic resistant tumor cells. Breast Cancer Res. 14, R2 (2012).

112. Ji, X. et al. Neutralization of TNFα in tumor with a novel nanobody potentiates paclitaxel-therapy and inhibits metastasis in breast cancer. Cancer Lett. 386, 24–34 (2017).

113. Kashimura, S. et al. Up-regulation of toll-like receptor-4 mRNA by docetaxel. Cancer Res. 64, 156–157 (2004).

114. Byrd-Leifer, C. A., Block, E. F., Takeda, K., Akira, S. & Ding, A. The role of MyD88 and TLR4 in the LPS-mimetic activity of Taxol. Eur. J. Immunol. 31, 2448–2457 (2001).

115. Resman, N. et al. Taxanes inhibit human TLR4 signaling by binding to MD-2. FEBS Lett. 582, 3929–3934 (2008).

116. Zimmer, S. M., Liu, J., Clayton, J. L., Stephens, D. S. & Snyder, J. P. Paclitaxel Binding to Human and Murine MD-2 *. J. Biol. Chem. 283, 27916–27926 (2008).

117. He, B. et al. Selective tubulin-binding drugs induce pericyte phenotype switching and anti-cancer immunity. EMBO Mol. Med. 17, 1071–1100 (2025).

118. Hattori, A. et al. Targeting Interleukin-6 as a Novel Strategy to Overcome Eribulin Resistance in Breast Cancer. Cancer Med. 14, e71273 (2025).

119. Dekkers, J. F. et al. Long-term culture, genetic manipulation and xenotransplantation of human normal and breast cancer organoids. Nat. Protoc. 16, 1936–1965 (2021).

120. Wells, C., Labban, N., Showalter, S. L., Przanowska, R. K. & Janes, K. A. Fast, flexible, learning-free organoid quantification and tracking with OrganoSeg2. BioRxiv Prepr. Serv. Biol.

121. Dowling, P. & Clynes, M. Conditioned media from cell lines: A complementary model to clinical specimens for the discovery of disease-specific biomarkers. PROTEOMICS 11, 794–804 (2011).

122. Runnebaum, I. B., Nagarajan, M., Bowman, M., Soto, D. & Sukumar, S. Mutations in p53 as potential molecular markers for human breast cancer. Proc. Natl. Acad. Sci. U. S. A. 88, 10657–10661 (1991).

123. Lehman, T. A. et al. p53 Mutations in human immortalized epithelial cell lines. Carcinogenesis 14, 833–839 (1993).

